# A compendium of *bona fide* reference markers for genuine plant extracellular vesicles and their degree of phylogenetic conservation

**DOI:** 10.1101/2024.11.21.624247

**Authors:** Miriam M. Rodríguez de Lope, Ibone R. Sánchez-Pajares, Estela Herranz, Cristina M. López-Vázquez, Ainara González-Moro, Alan Rivera-Tenorio, Carlos González-Sanz, Soledad Sacristán, Eduardo Chicano-Gálvez, Fernando de la Cuesta

## Abstract

Although the field of plant EVs (PEVs) is experiencing exponential growth, rigorous characterisation complying with MISEV guidelines has not been yet implemented due to the lack of *bona fide* reference markers. In this work, we have paved the way for the standardisation of PEV markers, providing the most profound proteomic data so far from apoplastic washing fluid-EVs, a sample enriched in genuine extracellular vesicles from plant tissue of two reference plant species: *Arabidopsis thaliana* (Arath-EVs) and *Brassica oleracea* (Braol-EVs). Besides, we analysed the protein content of the soluble fraction of the apoplast and calculated the enrichment of the potential markers studied in EVs. Additionally, we have conducted an exhaustive analysis of the proteomic data available so far from genuine EVs from any plant species, evaluating current potential markers, together with those found in our proteomic analyses. Our results provide evidence supporting the potential use of the following families as PEV markers: aquaporins, vacuolar-type ATPase complex subunits, some fasciclin-like arabinogalactan proteins (FLAs), tetraspanins, syntaxins, germin-like proteins and calreticulins. Next, we analysed the presence of orthologues and their degree of conservation throughout plant taxa, as well as in 2 reference species from the animal kingdom: human and mouse. Their degree of conservation was compared with that of current animal EV: CD63, CD81 and CD9. Among the protein families with potential to be used as PEV markers, 2 were found to be plant-specific: FLAs and germin-like proteins. On the other hand, aquaporins and vacuolar-type ATPase complex subunits showed the greatest degree of conservation across plant and animal kingdoms.

Our results provide key insights on several aspects of classical and novel protein identity markers for PEVs to assist in the selection of the best candidates for standardisation: 1) species-specific abundance, 2) specificity for PEVs, and 3) conservation and plant specificity.

## Introduction

Extracellular vesicles (EVs) from mammalian cell origin have been widely characterised and exhaustive guidelines to follow have been defined by the International Society for Extracellular Vesicles (ISEV), the so-called “Minimal Information for Studies of Extracellular Vesicles (MISEV)”, its more recent version being published in 2023^1^. Regardless of a constant and exponential growth of publications in the field of plant-derived EVs (PEVs), to date there is a lack of standardization in this particular scientific area. In fact, a call for rigor in the field of PEVs was made by Pinedo *et al.* in 2021 in the journal of the ISEV^2^, with a special focus on the need for specific markers that could allow to comply with the golden rules of EV characterization. Furthermore, ISEV has recently launched the plant EVs task force to gather experts in the field of PEVs to put forward initiatives regarding nomenclature, isolation, characterization, and standardization (https://www.isev.org/plant-task-force).

To date, several publications have provided data on PEVś cargo and potential markers, but a consensus on *bona fide* markers of PEVs has not yet been reached and thus, a response to the abovementioned call for rigor is still awaiting. One particular hurdle in this aim is the fact that most of the work in this direction has been performed in *Arabidopsis thaliana* L., a eudicot plant that stands as a reference model in the field, but there is not enough evidence supporting that these markers are applicable to plant EVs from other origin, especially monocots, bryophytes and algae. Studies in *A. thaliana* frequently use two tetraspanins, a protein group well-known to be involved in EV biogenesis and widely used as EV markers, to characterise and monitor PEVs: tetraspanin 8 (TET8) and tetraspanin 9 (TET9), the former being the most utilised. Another marker arising from *A. thaliana.* is PEN1, also called syntaxin of plants 121 (SY121), a member of the t-SNARE protein superfamily, which is related to the exocytic pathway^3^ and whose ortholog in humans, syntaxin-2, has been associated with exocytosis in mammalian cells^4^. This protein was also shown to be present in PEVs from *Nicotiana benthamiana*^5^. Nonetheless, it is unclear whether PEN1 is a marker of all types of EVs or of a specific subcategory^6^. On the other hand, a subset of EVs arising from a characteristic plant biogenesis pathway, exocyst-positive organelle (EXPO)-derived EVs, described in *A. thaliana* and *Nicotiana tabacum,* express the specific marker Exocyst complex component EXO70E2^7^. All these markers have mainly proven to be useful for studying PEVs from *A. thaliana* but translation into other species has not so far been robustly demonstrated.

One strategy for defining biomarkers of PEVs is the search for orthologues of mammalian EV markers. This has partially driven most of the abovementioned findings in *A. thaliana.*, with tetraspanins TET8 and TET9 being structurally similar to mammalian tetraspanin CD63^8^, with which they share less than 30% homology. Many other markers belonging to protein families related to EV biogenesis have also been explored: orthologues of Tumour susceptibility gene 101 protein (TSG101), ALG-2 interacting protein X (Alix), and Heat shock 70 kDa protein (Hsp70) have been found to be markers of EVs derived from plants in some studies. Regarding TSG101, called protein ELC-like in *A. thaliana*^9^, a homolog in lemon (*Citrus limon* (L.) Burm. f) juice-derived PEVs has been reported^6,10^. The ortholog of Alix in *A. thaliana,* vacuolar-sorting protein BRO1, has been proven to be involved in the biogenesis of vacuole and multivesicular bodies (MVBs)^11^. Besides, an Alix homolog has also been described in kiwi (*Actinidia chinensis Planch*.) pollen-derived PEVs, detected with an antibody targeting mammalian Alix^12^. Finally, orthologues of HSP70 have been found in EVs from tangerine^13^ and other citrus species^14^, as well as in the edible algae *Ecklonia cava* Kjellman^15^, the latter also detected using an antibody developed with the human protein as immunogen.

Therefore, there is an urgent need for a consensus in the field of PEVs to define the validity of current EV-specific markers, considering specificity, experimental evidence of abundance in EV isolates and conservation throughout the plant kingdom, in order to define their potential use for all plant sources or restricted to certain species. Concerning this, although systematic revision of all the available data in the field is crucial to answer this question, the lack of reproducible and robust data deposited in specialised repositories limits the potential of such approach. Hence, new experimental data may be required to deepen in the proteome of PEVs, aiming to find novel specific EV markers for plant vesicles and an important level of conservation in the plant kingdom that may allow their use for most, if not all, plant sources. With this goal in mind, we have conducted an in-depth proteomic analysis of genuine plant extracellular vesicles, achieved through the use of apoplastic washing fluid from plant tissue, from two well-annotated reference plant species: *A. thaliana* and *Brassica oleracea* L. Besides, we analysed the protein content of the soluble fraction of the apoplast and calculated the enrichment of the potential markers studied in EVs. Next, we exhaustively analysed the proteomic data published in the field, comparing them with the proteomes obtained in the present study. Our results allow us to provide more rigor and define the utility of the identity markers described so far, as well as defining novel PEV markers, by adding key data on EV and species specificity.

## Materials and Methods

### Plant Material

*A. thaliana* ecotype Columbia-0 (Col-0) seeds were sown, stratified at 4°C for 3 days in darkness, and moved to a growth chamber at 22°C, 80% relative humidity, under short day photoperiod (10-h light/14-h dark) and light intensity of 110-120 µE/m2 /s for 5 weeks. To obtain three biological replicates, plants were grown at different times under the same controlled conditions. Once they reached the same developmental stage, three independent batches were harvested.

As an alternative to *Arabidopsis* introducing a greater degree of variability to compare the robustness of the potential markers, we searched for a plant source grown in uncontrolled conditions, from a species well-annotated in proteomic databases, that being *Brassica oleracea*. Hence, fresh red cabbage (*Brassica oleracea var. capitata F. rubra*) purchased from a local market was used. Three biological replicates were obtained by selecting three whole cabbages, acquired on different days from the same market, ensuring they belonged to different production batches.

### Collection of apoplastic washing fluid (AWF) by vacuum infiltration

*A. thaliana* rosettes and cabbage leaves were processed separately to prevent cross-contamination and maintain biological independence. They were collected and thoroughly washed, first in 70% ethanol for surface sterilization and then rinsed twice with PBS to remove any residual ethanol. Due to the larger size of cabbage leaves, these were previously cut and washed thoroughly with PBS to avoid contamination with intracellular vesicles. The washed leaves were placed into a 100 mL syringe filled with PBS, and a gentle vacuum was applied to infiltrate the leaves. After the infiltration, the leaves were carefully positioned inside a 5mL plastic pipette tip, which was sealed with parafilm to keep the leaves in place. This setup was then inserted into a syringe without the plunger. The syringe was placed into a 50 mL Falcon tube, ensuring that the leaves stayed in place and did not touch the bottom. The tube was centrifuged for 15 min at 4°C at 1,000 x g to collect the AWF **(Fig 1)**.

**Figure 1.**
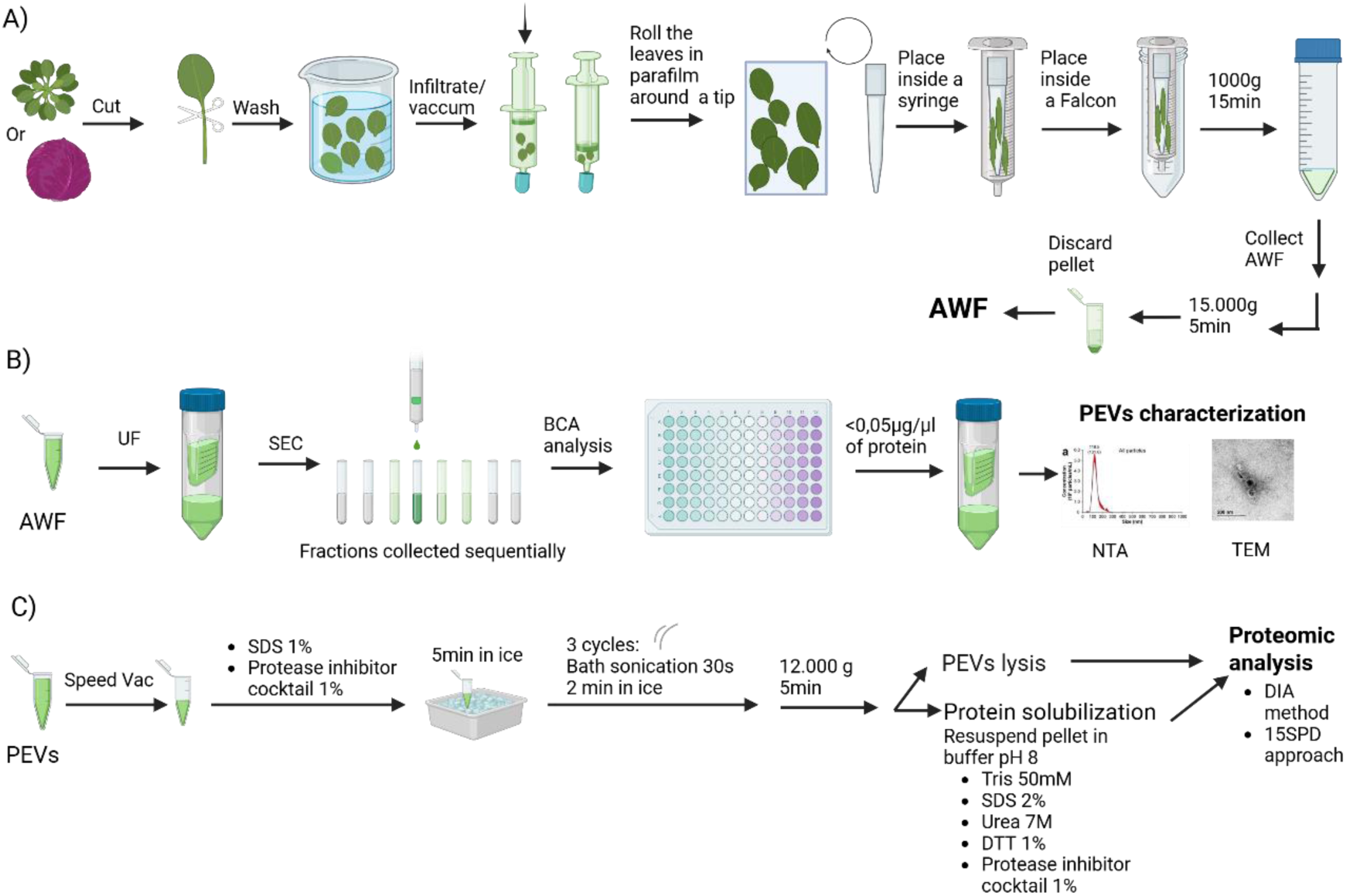
Workflow for Apoplastic Washing Fluid collection, plant extracellular vesicle isolation, and sample preparation for proteomics. A) Apoplastic Washing Fluid (AWF) extraction from A. thaliana or B. oleracea. B) Plant EVs (PEVs) isolation using ultra filtration (UF) and size exclusion chromatography (SEC). C) PEVs lysis and sample preparation for proteomics based on a two-step solubilisation.

To assess the dilution factor of the recovered AWF a parallel infiltration was performed in *A. thaliana* using a 50 µM indigo carmine solution, as described elsewhere^16^. Indigo carmine was added to the infiltration buffer (PBS) and leaves were infiltrated as above. Leaves were then washed in distilled water to remove excess dye from the surface before centrifugation. Absorbance at 610 nm (OD₆₁₀) was measured for both the infiltration solution and the recovered AWF. The apoplast dilution factor was calculated as:

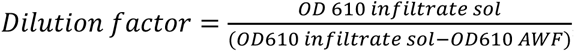, being OD_610_ infiltrate sol the absorbance of the 50 µM indigo carmine solution.

Cytosolic contamination of the apoplastic fluid was assessed by measuring the activity of the enzyme malate dehydrogenase (MDH), following the method described by van den Dries N. *et al.*^17^ with slight modifications. Briefly, the reaction mixture was prepared in a final volume of 250 µL, containing 125 µL of Tris-HCl buffer (0.1 M, pH 7.8), 50 µL of distilled water, 25 µL of 1 mM oxaloacetic acid, and 25 µL of AWF. The reaction was initiated by adding 25 µL of 0.4 mM NADH. The decrease in absorbance at 340 nm, corresponding to NADH oxidation, was monitored using a microplate reader for 5 minutes. To determine total MDH activity in whole-leaf homogenates, leaf tissues were ground in liquid nitrogen, and proteins were extracted using a buffer composed of 50 mM Tris-HCl (pH 7.5), 200 mM NaCl, 1 mM EDTA, 10 mM NaF, 2 mM sodium orthovanadate, 1 mM sodium molybdate, 10% (v/v) glycerol, 0.1% (v/v) Tween-20, 1 mM dithiothreitol (DTT), and 1 mM phenylmethylsulfonyl fluoride (PMSF). Samples were centrifuged at 20,000 × g for 30 minutes at 4 °C, and the supernatant was used for the MDH assay as described above instead of the AWF. The MDH activity detected in the apoplastic fluid was expressed as a percentage of the total MDH activity measured in the leaf homogenate.

### Isolation of PEVs from AWF

The AWF was centrifuged for 5 min at 4°C at 15,000 x g to remove cell debris and then filtered through a 0.22 μm filter (Jet Biofil). Next, the supernatants (the clean AWF) were concentrated by ultrafiltration using Amicon Ultra-15 100 kDa filters (Millipore, Merck) at 3500 x g at 4°C. The concentrated supernatants were then subjected to size exclusion chromatography (SEC) using in-house casted Sepharose CL-2B (Sigma-Aldrich Chemical Co) columns to isolate EVs. Protein quantification of the fractions collected from SEC was performed using the Pierce BCA method (Thermo Fisher Scientific) and fractions containing less than 0,05μg/μL of total soluble protein were further concentrated by ultrafiltration with Amicon Ultra-15 10 kDa filters (Millipore, Merck) at 3,500 x g at 4°C. The isolation and characterization process are illustrated in **Figure 1**.

### Characterization of PEVs by nano tracking analysis (NTA) and transmission electron microscopy (TEM)

EVs isolated from *A. thaliana* (Arath-EVs) and *B. oleracea* (Braol-EVs) AWF were characterised by nanoparticle tracking analysis (NTA), using a NanoSight LM10 instrument coupled to an automated Syringe Pump module (Malvern Panalytical), measuring their concentration and size distribution. Five technical measurements of each biological replicate were performed. The same camera level, threshold, and focus were used for all measurements.

EVs morphological characteristics were observed by transmission electron microscopy (TEM). For this purpose, they were fixed in 0.1 % PFA. A 15 μL droplet of sample was placed in an electron microscopy Formvar/carbon grid (Ted Pella) for 10 min. After 5 min in bidistilled water, a negative staining was conducted using 20 μL of uranyl acetate 2% for 30 s. The samples were visualized in a JEM 1010 (JEOL Peabody) at the Microscopy Unit, Faculty of Medicine UAM.

### Proteomic analysis of PEVs

#### Sample preparation

EV samples (1×10^10^ particles) were concentrated using a SpeedVac (Thermo Scientific) up to an approximate volume of 50 μl. Samples were then lysed by adding an EV lysis buffer: SDS 1% (Sigma-Aldrich) and a protease inhibitor cocktail 1% (Sigma-Aldrich). The mixture was placed 5 min on ice, followed by 3 cycles of sonication in a water bath sonicator (Fisher Scientific) with each cycle consisting of 30 s of sonication followed by 2 min on an ice bath.

The samples were then centrifugated for 5 min at 4°C at 12,000 x g and proteins in the resulting pellet were solubilized with a protein solubilisation buffer (50mM Tris, SDS 2%, 7M Urea, DTT 1%, protease inhibitor cocktail 1%, pH 8,0). The amount of protein in both extracts was quantified by BCA, and samples were sent to the IMIBIC Mass Spectrometry and Molecular Imaging Unit for further analyses (**Fig. 1C**). Total protein was further quantified in the Mass Spectrometry Unit using the Qubit Protein Assay Kit (Thermo Fisher Scientific, Waltham, MA, USA) according to the manufacturer’s instructions. The samples were then digested using the iST kit (PreOmics, Germany) and peptides were diluted to 10 ng/µl in LC-MS Water containing 0.1% (v/v) formic acid. A pooled sample was prepared by combining equal volumes of each individual sample. This pooled sample was used exclusively for intra-experimental quality control to ensure consistent acquisition performance throughout the experiment. Subsequently, 400 ng of the peptide solution were loaded onto Evotips (Evosep) following the manufacturer’s instructions. Three biological replicates (n=3) were prepared for analysis of samples coming from both *A. thaliana.* and *B. oleracea.* In addition, Pierce™ Hela Tryptic Digest Standard (Thermo Scientific) was used to equilibrate and activate the LC system. These digests were spiked with iRT peptides (Biognosys, Switzerland) and monitored in real time (mass deviation, ion mobility, retention time) using ProteoScape software (Bruker) to ensure optimal instrument performance before sample acquisition. Three technical replicates from each sample were analysed using Liquid chromatography–mass spectrometry (LC-MS/MS) in Data-independent acquisition and parallel accumulation—serial fragmentation (DIA-PASEF) mode on the Evosep One TIMSTOF-Flex Clinical Proteomics Platform of the Maimonides Institute of Biomedical Research of Cordoba (IMIBIC) Mass Spectrometry and Molecular Imaging Unit (IMSMI) as follows: <u>Liquid chromatography</u>

LC-purified tryptic digests were separated using the preset 15 SPD method (88-min gradient time, 400 ng peptides) with an Evosep One LC system. A 10-μm fused silica ID emitter (Bruker Daltonics) was inserted into a nanoelectrospray source (Bruker Daltonics CaptiveSpray). The emitter was linked to a 15-cm × 150-μm reverse-phase column with 1.9-μm C18 beads (PepSep Fifteen columns, Bruker Daltonics), and column was heated to 60 °C in an oven compartment (Bruker Daltonics). The mobile phases, water and acetonitrile, were buffered with 0.1% formic acid (LC-MS grade, Fisher Scientific).

#### Mass Spectrometry

LC was connected online to a TIMS Q-TOF apparatus (timsTOF Flex, Bruker Daltonics) via a CaptiveSpray nano-electrospray ion source and DIA-PASEF methods were used^18^. Ion mobility dimension calibration was done with 3 Agilent ESI-L Tuning Mix ions (m/z, 1/K0: 622.0289 Th, 0.9848 Vs cm−2; 922.0097 Th, 1.1895 Vs cm−2; 1221.9906 Th, 1.3820 Vs cm−2). Collision energy was linearly decreased from 59 eV at 1/K0 = 1.6 Vs cm−2 to 20 eV at 1/K0 = 0.6 Vs cm−2. In addition, a ‘long gradient’ method was applied (m/z range: 400–1201.0 Th, 1/K0 range: 0.6– 1.6 Vs cm−2, diaPASEF windows: 16 × 25 Th).

#### Data Analysis

Data identification and quantification were conducted using Spectronaut proprietary software (https://biognosys.com/software/spectronaut/). A directDIA approach (Spectronaut 18, Biognosys) analysis was done using the entire sample dataset and quality-control (QC) runs to monitor system performance. By using UniProt (https://www.uniprot.org/) as reference database, a subset of proteins formatted in fasta file format was built for *A. thaliana* (Taxon 3702) (September, 2023) and *B. oleracea* (Taxon 109376) (May, 2024). A Global data normalization of label free-quantitation (LFQ) parameter media was performed with Spectronaut18, followed by non-assigned numbers (NaN) data imputation by 1/5 limit of detection (LOD) using Metaboanalyst^19^. After normalization protein intensities were filtered by abundance. To calculate enrichment ratios, the mean of normalized LFQ values was calculated for each condition, and the enrichment degree was obtained as the ratio = LFQ PEVs/ LFQ soluble fraction.

The raw proteomics data and the minimum information about a proteomics experiment (MIAPE), complying with the Proteomics Standards Initiative (PSI) of the Human Proteome Organization (HuPO) have been deposited to the ProteomeXchange Consortium via the PRIDE^20^ partner repository with the dataset identifier PXD058098.

### Enrichment analysis

Protein-protein interactions were first analysed using STRING^21^. Default parameters were applied, using “Medium confidence (0.400)” as the minimum required interaction score and hiding the unconnected nodes. “Functional enrichment visualization” graph was obtained applying default parameters defined by STRING. To perform a specific enrichment analysis of the list of proteins interacting inside our group of interest selected from the “Functional enrichments”, this protein-protein interaction network was downloaded, applying the default parameters except in the case of the minimum required interaction score, which was increased to “High confidence (0.700)”. The proteins from this network were implemented in Cytoscape^22^, where we employed the “String enrichment” application to perform a “Retrieve functional enrichment”, followed by filtering of concepts by the category “GO Cellular Component”, from which an Enrichment Map was created by applying a “Connectivity cut-off” of 0.4.

### Comparison with other proteomes

To compare the proteomic entries from Arath-EVs with the 3 reference datasets obtained from publications of *A. thaliana* AWF-derived EVs, for some datasets, we transformed NCBI IDs to UniProt IDs and we edited the results from our list of Arath-EVs, since more than one protein could be potentially identified in some cases for a single entry. Therefore, the list was modified with a text editor by separating the proteins that appeared together in the same entry to consider them individually, resulting in a total number of 1445 protein identifiers. This list of proteins was then confronted with the 3 reference datasets, and a Venn diagram was done in jvenn software (https://jvenn.toulouse.inrae.fr/app/example.html) to obtain a final list of common proteins.

BLASTP tool of Uniprot was used to compare the proteomic entries from Arath-EVs with the 7 datasets obtained from publications of different PEVs to verify the presence of orthologues in other proteomes. For most species, Uniprot was used for this comparison; however, in the case of sunflower, a specific database was used (https://www.heliagene.org/HanXRQ-SUNRISE/) due to the lack of comprehensive sunflower entries in Uniprot.

### Orthology search

The identification of orthologues for the proteins of interest was performed using the BLASTP tool of UniProt. For proteins whose orthology was studied primarily within plant kingdom phylogenies, *A. thaliana.* FASTA sequences were queried against the species listed in **Table 1A**. All sequences were obtained from UniProt, except for *Picea abies* (<u>L.</u>) H. Karst, which is not well-annotated in this database. Therefore, we obtained its protein sequences and identity percentages using the Evorepro database and the BLAST tool provided by this resource (https://evorepro.sbs.ntu.edu.sg/blast/).

**Table 1.** Species used for orthology search. A) Species for orthology search of potential PEVs markers, B) Species for orthology search of well-defined animal EVs markers.

| SPECIES | PHYLOGENY | MNEMONIC NAME | TAXON ID |
| --- | --- | --- | --- |
| A |  |  |  |
| **Arabidopsis thaliana** | Plantae, angiosperms, eudicots | ARATH | 3702 |
| **Brassica oleracea var. oleracea** | Plantae, angiosperms, eudicots | BRAOL | 109376 |
| **Oryza sativa subsp. japonica** | Plantae, angiosperms, monocots | ORYSJ | 39947 |
| **Solanum lycopersicum** | Plantae, angiosperms, eudicots | SOLLC | 4081 |
| **Picea abies** | Plantae, gymnosperms, pinopsida | PICAB | 3329 |
| **Marchantia polymorpha (Common liverwort)** | Plantae, marchantiophyta, marchantiopsida (bryophyte) | MARPO | 3197 |
| **Chlamydomonas reinhardtii** | Plantae, chlorophyta, chlorophyceae | CHLRE | 3055 |
| **Homo sapiens** | Animalia, chordata, mammalia | HUMAN | 9606 |
| **Mus musculus** | Animalia, chordata, mammalia | MOUSE | 10090 |
| B |  |  |  |
| **Homo sapiens** | Animalia, chordata, mammalia | HUMAN | 9606 |
| **Pan troglodytes** | Animalia, chordata, mammalia | PANTR | 9598 |
| **Mus musculus** | Animalia, chordata, mammalia | MOUSE | 10090 |
| **Rattus norvegicus** | Animalia, chordata, mammalia | RAT | 10116 |
| **Oryctolagus cuniculus** | Animalia, chordata, mammalia | RABIT | 9986 |
| **Gallus gallus** | Animalia, chordata, birds | CHICK | 9031 |
| **Xenopus laevis** | Animalia, chordata, amphibia | XENLA | 8355 |
| **Danio rerio** | Animalia, chordata, actinopterygii | DANRE | 7955 |
| **Drosophila melanogaster** | Animalia, arthropoda, insecta | DROME | 7227 |
| **Caenorhabditis elegans** | Animalia, nematoda, secernentea | CAEEL | 6239 |

For proteins defined as EV markers in animals, *H. sapiens* FASTA sequences were queried against the species listed in **Table 1B**.

For each protein of interest, we identified the putative orthologues from each species with the highest identity score according to the UniProt classification. MEGA 11 software^23^ was used to construct phylogenetic trees from the protein sequences found for all considered species. First, the alignment of the FASTA files for each group of putative orthologues was performed with the ClustalW tool and then, phylogenetic trees were generated with the “Neighbour-joining” method.

## Results

### AWF Yield and Purity

Prior to infiltration, the fresh weight of *A. thaliana* leaves averaged 9.5g (35 plants), with a recovery of 0.26 mL of AWF per gram of fresh weight. A lower yield was obtained with *B. oleracea*, from which 130.4g of fresh tissue (corresponding to half red cabbage) yielded a recovery of 0.092 mL/g of fresh tissue.

Due to the fact that intense pigmentation of red cabbage interferes with the wavelengths required to measure indigo carmine dilution and MDH activity, these analyses were performed only using *A. thaliana* AWF, as a proof-of-concept.

Based on the dilution of indigo carmine, the calculated dilution factor for *A. thaliana* AWF was 2,26 ± 0,76. To confirm that the extracted apoplastic fluid was originated from the apoplast, cytoplasmic contamination was evaluated by measuring malate dehydrogenase (MDH) activity. The contamination level was 9.1% relative to total MDH activity, calculated using *A. thaliana* leaf lysate. This value is comparable to the ∼10% contamination reported by van den Dries N. et al.^17^ at 3000 × g for *A. thaliana* AWF. Moreover, it is lower than the level reported by O’Leary B.M. et al.^16^ for *Phaseolus vulgaris* AWF, which was 2.5 ± 0.9 U/mL; in our case, the equivalent value is 0.58 ± 0.009 U/mL.

### Isolation and Characterization of Arath-EVs and Braol-EVs

Vesicles were successfully extracted from the AWF of *A. thaliana* and *B. oleracea* leaves. After size-exclusion chromatography, eleven 1mL fractions were collected for *A. thaliana* and thirteen 1mL fractions for *B. oleracea*, being these the fractions with a soluble protein content below 0,05μg/μL. Three batches of each plant species were purified, and the concentration and size distribution were observed by NTA (**Fig. 2A**). PEVs from both plant species had a similar size, being the mean for Arath-EVs of 157,82 ± 7,35 nm, and the mean for Braol-EVs 159,13 ± 4,35 nm (**Fig. 2B**). However, yield obtained from Arath-EVs was 1,61 x 10^9^ ± 5,61x 10^8^, higher than yield from Braol-EVs, 8,57 x 10^7^ ± 6,03 x 10^7^ (**Fig. 2C**). In all the samples assayed, TEM demonstrated a typical spherical vesicular appearance, with EVs showing integrity and the expected size (**Fig. 2D**).

**Figure 2.**
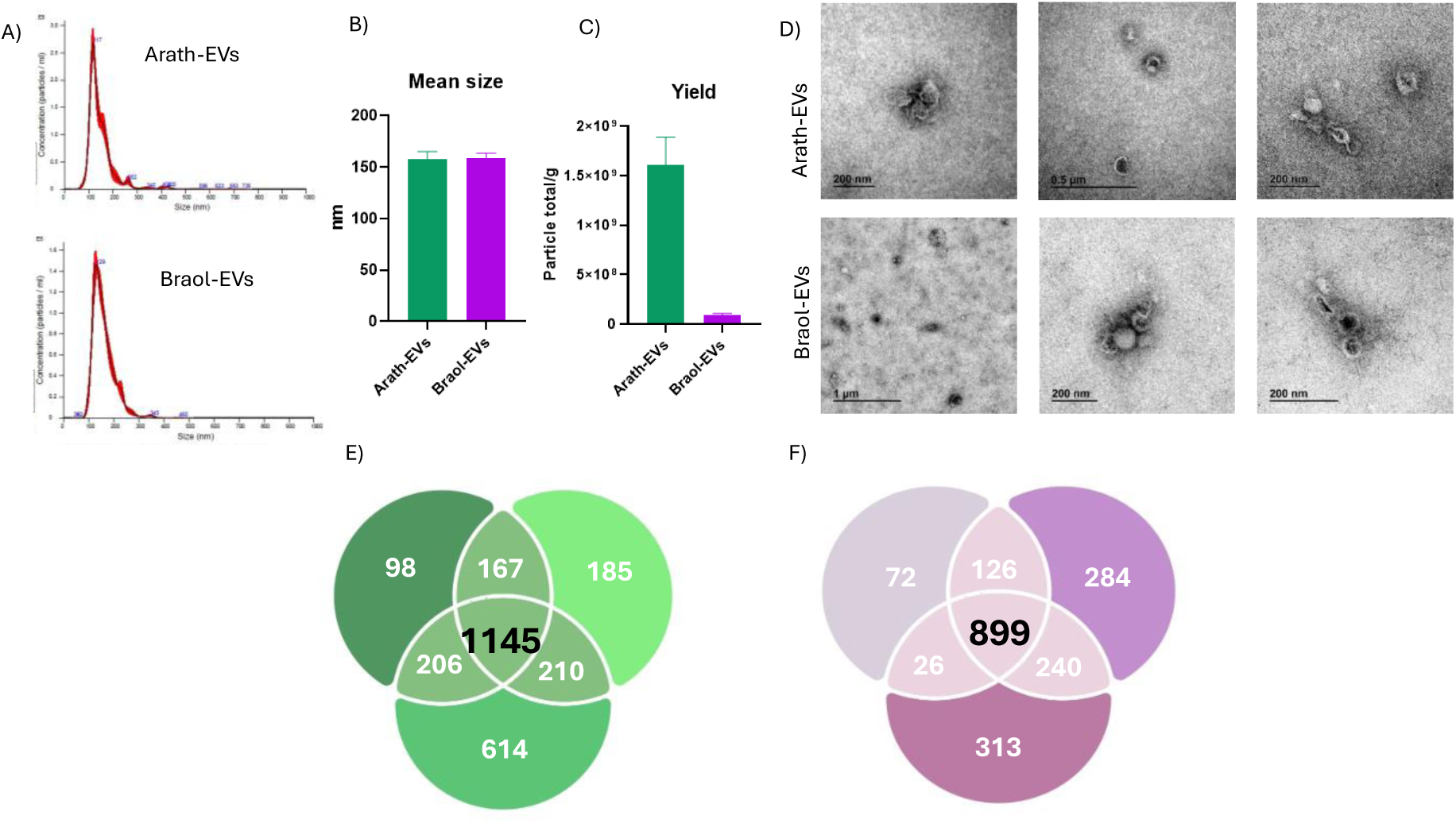
Characterization of A. thaliana and B. oleracea PEVs and proteomic analysis. A) NTA profile of Arath-EVs and Braol-EVs extracted from apoplastic washing fluid B) Calculated mean size the three batches of PEVs extracted. C) Yield of the three batches of PEVs extracted D) TEM analysis of the PEVs. E) Venn diagram of proteins found in each of the three replicates after protein solubilisation of Arath-EVs. E)Venn diagram of proteins found in each of the three replicates of the extract after protein solubilisation from Braol-EVs.

### Analysis of the proteome of PEVs

In order to extract proteins from PEVs, we optimized the extraction protocol for SEC-isolated EVs combined with LC-MS/MS, based on a two-step solubilisation. As part of the optimization, extracts from Arath-EVs coming from both solubilisation steps were analysed separately by proteomics to inspect which one had more proteins as well as to check their complementarity. Three biological replicates of both samples were then analysed and Venn diagrams were built up in order to focus only on the proteins that were present in all 3 replicates. After the EV lysis (first step) 117 proteins were identified **(Suppl. Fig. 1A)**, compared to 1145 proteins identified after the protein solubilisation **(Fig. 2E)** (second step). An amount of 94 proteins were identified in common, with only 23 proteins identified only in the first step (**Suppl. Fig. 1B**). The proteins identified in each replicate are listed in the **Suppl. Table 1**, organized by abundance, calculated using their normalized LFQ; including also the 23 proteins only identified in the first step. The principal component analysis (PCA) of both samples is shown in **Suppl. Fig. 2A.**

From this moment on, we decided to analyse only the extract after the second step of solubilisation in Braol-EVs samples. Three biological replicates were analysed and 899 proteins were found in common for Braol-EVs (**Fig. 2F**). The proteins identified are listed in the **Suppl. Table 2**, organized by abundance, calculated using their normalized LFQ.

Protein-protein interaction analysis was performed to narrow down the networks of the most relevant proteins with potential utility as PEV markers. To do so, we analysed by the software STRING the list of 1145 proteins from the Arath-EVs proteomics. We reviewed the “Functional enrichment visualization” graphs generated and, specifically, the one belonging to the “Cellular component” category, which showed the term “Secretory vesicle” occupying the fifth position in the top 10 enriched terms (**Fig. 3A**). Hence, we decided to perform a specific functional enrichment analysis with Cytoscape of the proteins within this term, to get a clearer picture of which other cellular compartments may be related to this group of proteins (**Fig. 3B**). In this network, the proximity of “Secretory vesicle” term to “Plant-type vacuole” and “Vacuole” terms is notable, aligning with the role of vacuoles as key players in the biogenesis of PEVs, suggesting they could serve as robust PEVs markers. Additionally, the “anchored component of membrane” term stood out, containing 6 fasciclin proteins (out of 9 proteins), also appearing in secretory vesicle and membrane terms. Next, we selected all proteins with greater potential as PEV markers across these four terminologies and generated a protein-protein interaction network using STRING to highlight these associations (**Fig. 3C**). Similarly, we performed the same enrichment analysis by the software STRING with the list of 899 proteins commonly found in Braol-EVs. In this case, the term “Secretory vesicle” also appeared among the top enriched terms, occupying the seventh position in the “Cellular Component” category **(Suppl. Fig. 3).**

**Figure 3.**
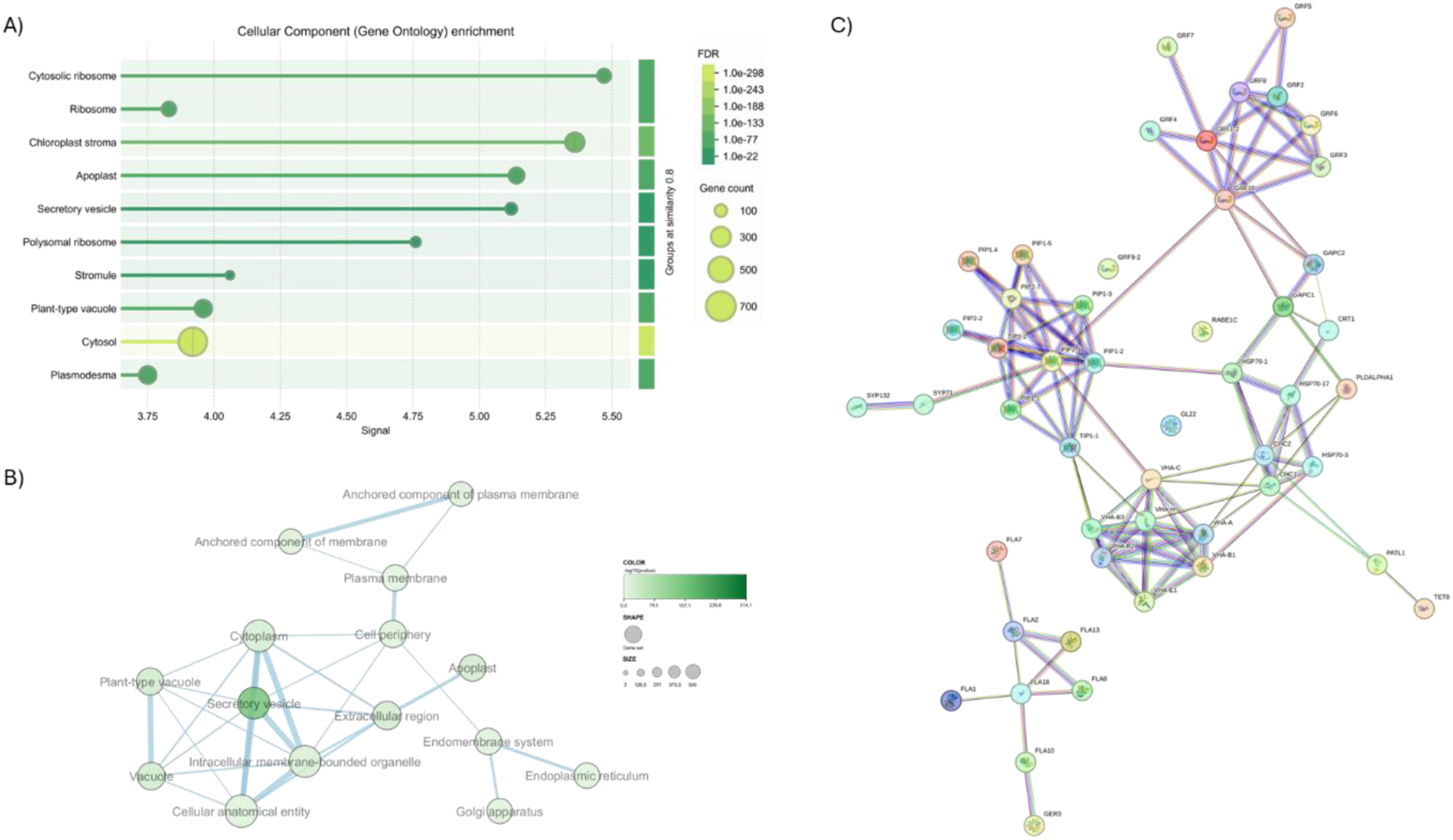
Protein-Protein Interaction and Functional Enrichment Analysis of Arath-EV Proteins. A) Cellular Component enrichment analysis of the 1145 proteins identified in 3 biological replicates of Arath-EVs. B) Network displaying specific functional enrichment for proteins within the “Secretory vesicle” term, revealing connections to related terms such as “Plant-type vacuole” and “Vacuole,” indicating shared biosynthetic pathways relevant to PEVs. C) Protein-Protein interaction network of potential PEV markers. Protein-Protein interactions were analysed using STRING ((https://string-db.org/) and implemented in Cytoscape^22^.

Focusing on the proteins obtained after this analysis, we searched for their orthologues in Braol-EVs, selecting the protein families identified in PEVs from both species. Among these, 12 families of proteins were chosen as potential markers present in both EVs: fasciclin-like arabinogalactan proteins (FLAs), germin-like protein, patellins, heat shock proteins, aquaporins, calreticulins, clathrins, phospholipase D, Rab proteins, Vacuolar-type ATPase (supp ATPase) proteins, 14-3-3-like proteins and glyceraldehyde-3-phosphate dehydrogenases (GADPHs) (**Table 2**).

**Table 2.**
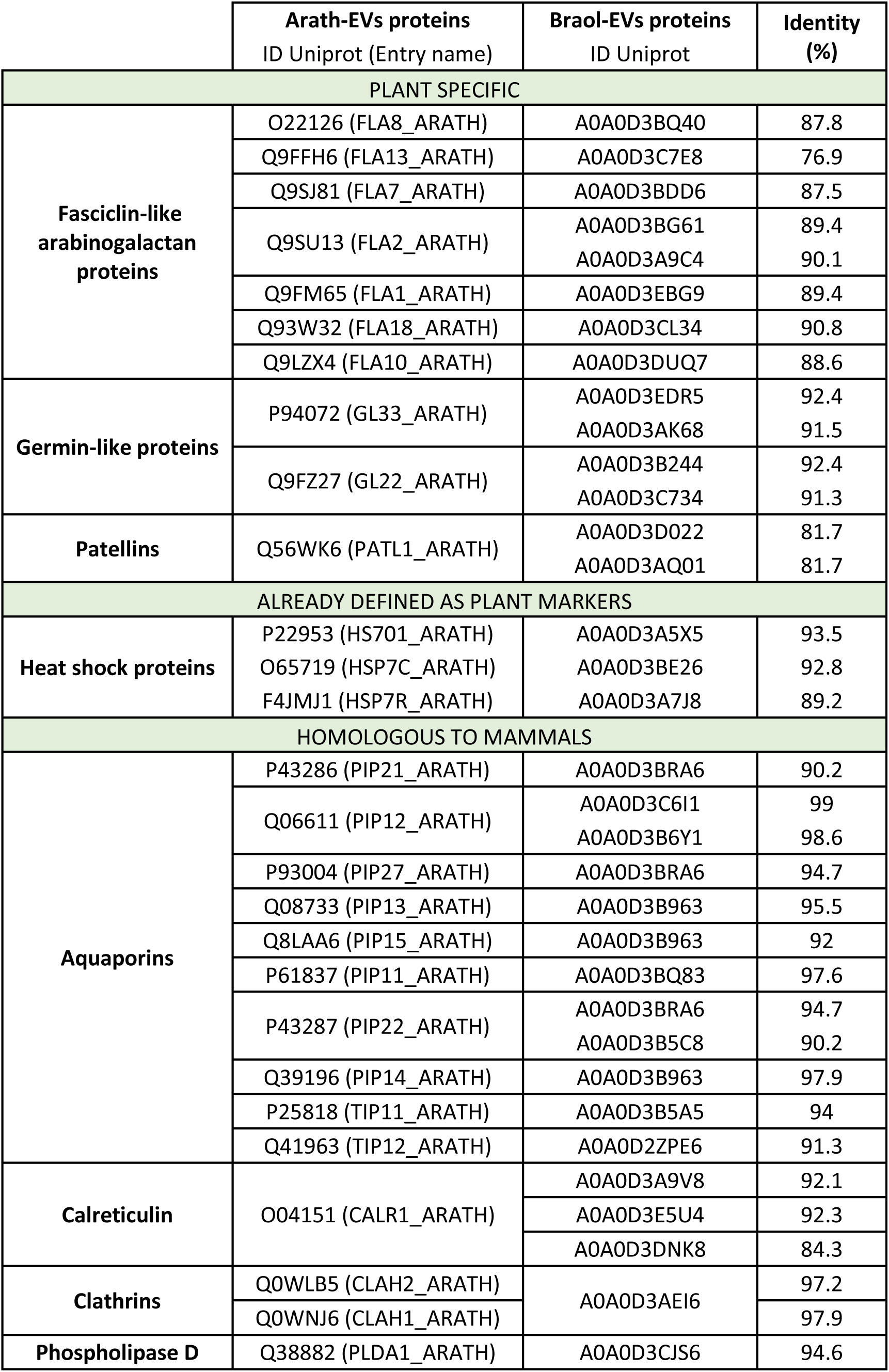
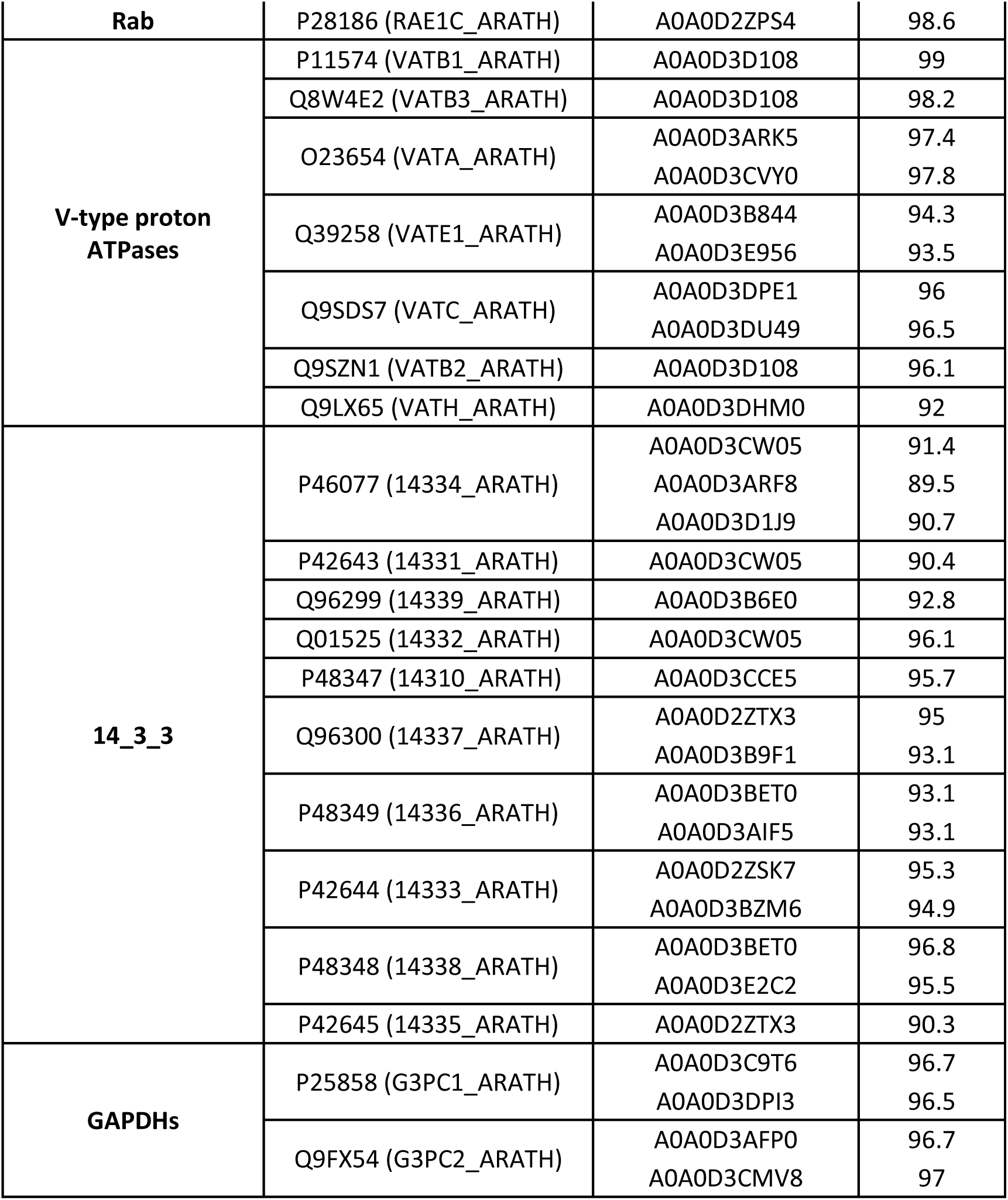
Putative protein markers. This table presents 12 protein families selected as potential markers due to their biological relevance.Proteins identified in both Arath-EVs and Braol-EVs are sorted by abundance calculated with the LFQ overall mean based on Arath-EVś protein list and their percentage of identity is displayed.

### Comparison with previously published PEV *A. thaliana* proteomes

The 12 protein families found in common on the proteomic results of Arath-EVs and Braol-EVs, were analysed by a second approach to explore their presence in other EVś proteomics studies.

As *A. thaliana* is the best annotated species in the UniProt database, we compared the proteomic data obtained in the present study with 3 reference datasets from publications of *A. thaliana* AWF-derived EVs^3,24,25^. The size of each list appears in **Figure 4A**. 34 proteins were found in common between the four lists (**Fig. 4B and C**). Of these 34 proteins, 27 appeared in our data of Braol-EVs (underlined in **Fig. 4C**). The UniProt IDs and % of identity of these proteins are shown in **Suppl. Table 3**. This list includes the previously mentioned FLAs, heat shock protein, aquaporins, clathrins, phospholipase D, Rab proteins, V-type proton ATPases and 14-3-3 family proteins. Additionally, some unspecific proteins with low evidence to be present in EVs were also identified, which were subsequently excluded from further analyses. Notably, one protein family of common EV-related proteins consistently identified on all Arath-EVs studies but not found in our previous analysis emerged: the syntaxins, which we could not identify in Braol-EVs.

**Figure 4.**
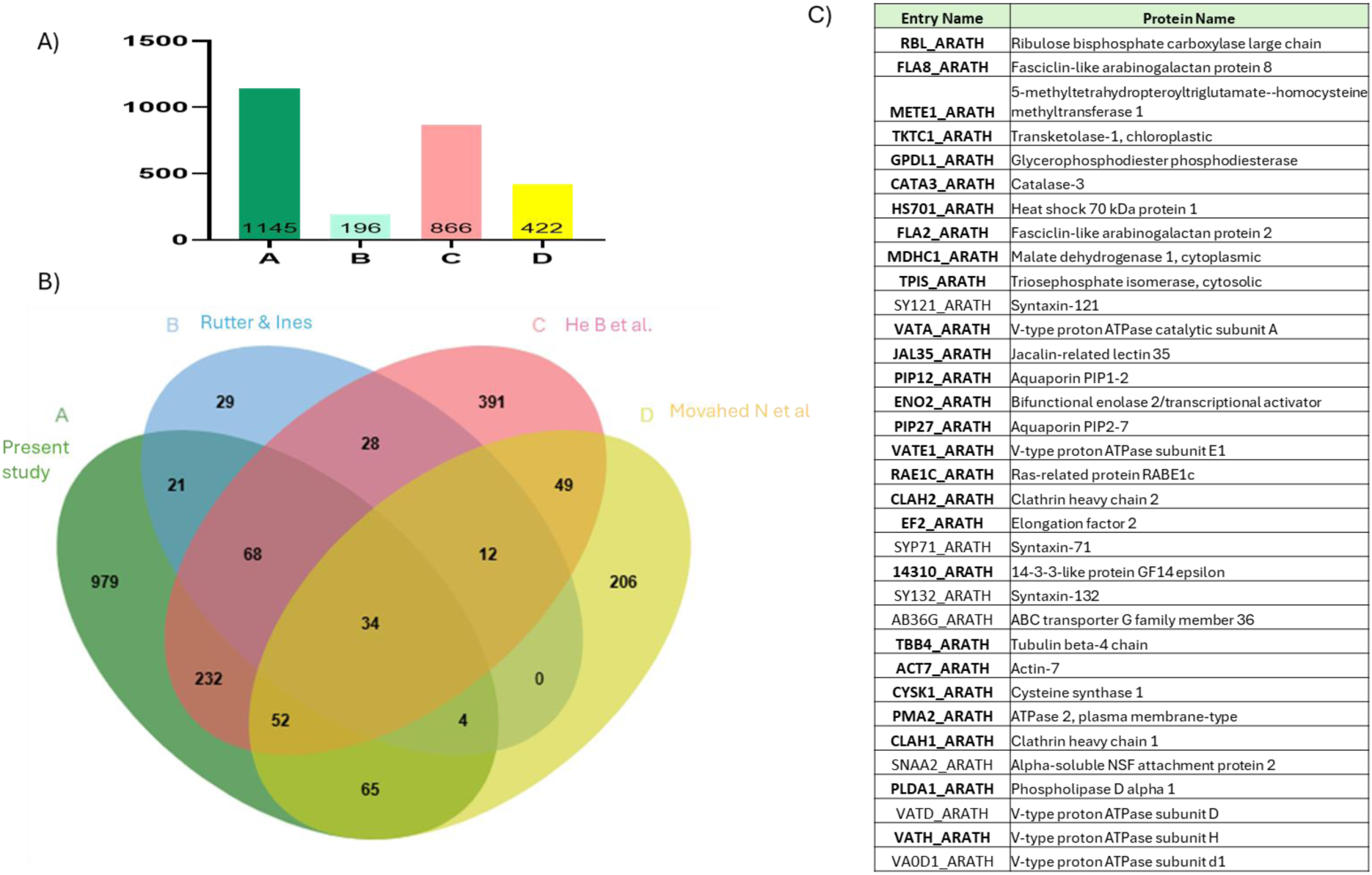
Comparative Analysis of A. thaliana vesicle proteins. A) Bar diagram showing the number of vesicle-associated proteins identified in each of the four studies used. B) Venn diagram illustrating the 34 proteins shared across all datasets. C) Table listing these 34 common proteins, with proteins also identified in our Braol-EVs dataset highlighted in bold.

To assess the potential of these candidates as global markers for PEVs, we generated a unified list of proteins shared by both approaches. This merged list included the 12 families shown in **Table 2** plus the syntaxin family, as identified in in **Figure 4C**. We also added the tetraspanin family, as TET8 is suggested as a PEVs marker in multiple studies^6^. TET8 was identified in our Arath*-*EVs proteomics data, but was absent in two out of the 3 Braol*-*EV replicates. Additionally, TET8 appeared in two out of the three *A. thaliana* reference studies previously published.

### Enrichment analysis

To evaluate the specificity of the highlighted proteins, we also submitted for proteomic analysis three replicates of the fractions containing more than 0,05μg/μL of total soluble protein after chromatography of Arab-EVs. These fractions are characterised by a higher content of soluble proteins and a lower concentration of vesicles compared to the EV-enriched fractions (fractions that we used to isolate the vesicles). As with the vesicle fractions, Venn diagrams were generated to identify proteins consistently present across all three replicates, resulting in 1,370 shared proteins **(Suppl. Fig. 1C).** By comparing the abundance of each protein in both types of fractions, we performed an enrichment analysis to evaluate whether the candidate proteins are specifically enriched in EVs, as showed in **Figure 5A** and in **Suppl. Table 4. Suppl Fig. 2B** shows the principal component analysis (PCAs) of both samples. Proteins that are either exclusively present in the vesicle-containing fractions or significantly enriched are ranked by their abundance in **Figure 5B**.

**Figure 5.**
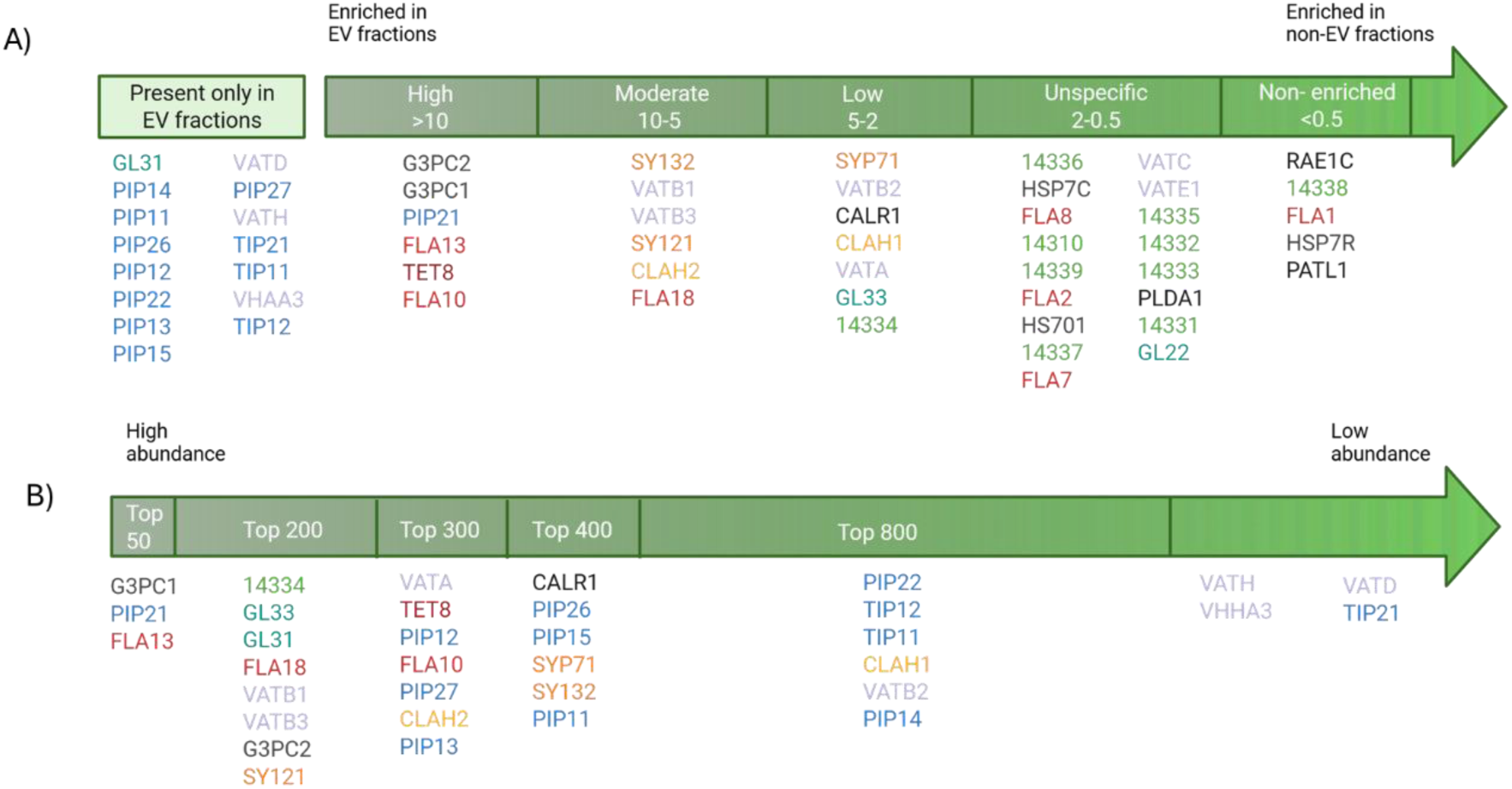
Identification of potential PEV marker proteins: protein enrichment in Arath-EVs and experimental abundance estimation. A) Proteins of interest ranked by their enrichment in EVs B) Enriched proteins ordered by their potential abundance, calculated using the normalized LFQ parameter in Arath-EVs.

### Comparison with previously published PEV proteomes

Finally, we conducted a systematic revision of all the proteomic studies on EVs from other plant sources to verify the frequency of these proteins as potential markers. The proteomic datasets selected had to fulfil the following criteria: 1) Isolation of true extracellular vesicles (intracellular vesicles underrepresented); 2) Depth of > 100 proteins identified; 3) Valid isolation methods according to MISEV guidelines. This search yielded 8 publications: 3 on AWF from sorghum^26^ and sunflower^27^, 3 on juice from pomegranate^28^ and diverse citrus plants^10,14^ 1 on callus secretome from pine^29^ and 1 on pollen secretome from kiwi^12^ **(Table 3).** All the protein families selected appear recurrently in other proteomic studies, which corroborates their presence in different sources of PEVs **(Table 4).**

**Table 3.** Cross -study comparation. Selected proteomic datasets from plant-derived EVs used in the comparison.

|  |  |  | Nº of proteins | Authors |
| --- | --- | --- | --- | --- |
| AWF | Shorgum | *Sorghum bicolor* | 437 | Chaya et al. (2024) |
|  | Sunflower | *Helianthus annuus* | 237 | Regente et al. (2017) |
| Polen | Kiwi | *Actinidia deliciosa* | 1170 | Suanno et al. (2023) |
| Callus | Pine | *Picea abies* | 557 | Kankaanpää et al. (2024) |
| Juice | Pomegranate | *Punica granatum* | 145 | Sánchez-López et al. (2022) |
|  | Citrics | *Citrus sinensis*, *C. limon*, *C. paradisi*, *C. aurantium* | 600-800 | Pocsfalvi et al; Raimondo et al. (2018, 2015) |

**Table 4.**
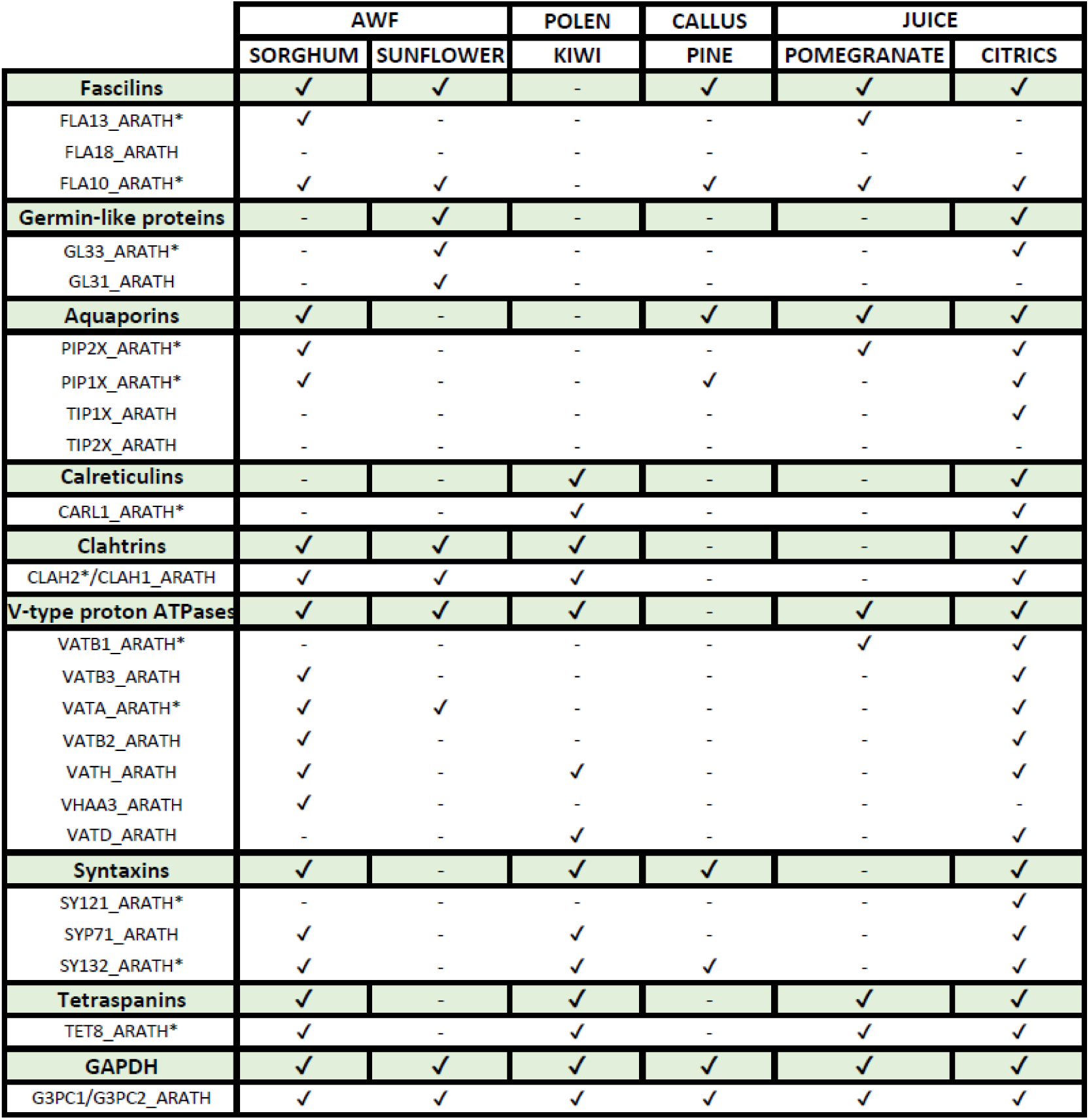
Presence / absence of selected protein families and family members across various PEVs. The table lists protein families and subfamilies, indicating the presence (✔) or absence (-) in the analysed species. High sequence similarity did not allow reliable differentiation between PIP1, PIP2, TIP1 and TIP2 isoforms by BLAST; subfamilies were thus analysed as groups. CHAH2/CLAH1 and G3PC1/G3PC2 were also grouped for this analysis due to their similar identity scores.

### Protein orthology and phylogenetic distance

Next, an orthologues′ search was conducted by BLASTP using UniProt, with *A. thaliana* protein sequences queried against those of six representative species from each subdivision of the plant kingdom, together with two mammalian species (human and mouse). All IDs chosen to make the phylogenetic trees appear in **Suppl. Table 5** We evaluated the conservation degree of these proteins among representative phylogenies, as well as their preservation between kingdoms. Furthermore, a similar analysis was conducted for animal CD63, CD81, and CD9, which are well established EV markers, providing a reference to help validate our proposed proteins as potential PEV markers.

To illustrate the evolutionary relationships among putative orthologues, phylogenetic trees were generated using MEGA11, with all trees represented using a uniform scale of 0.05. Phylogenetic trees for a selection of proteins listed in **Table 3** (labelled with *) Care provided as supplementary materials. In **Figure 6**, representative trees for some proteins from certain specific families are shown. Since more abundant proteins could represent potential markers, we removed the less abundant proteins, specifically those ranked from position 800 onward according to **Fig. 5B**. 14334 was also removed since all the others 14-3-3 proteins were found to be unspecific.

**Figure 6.**
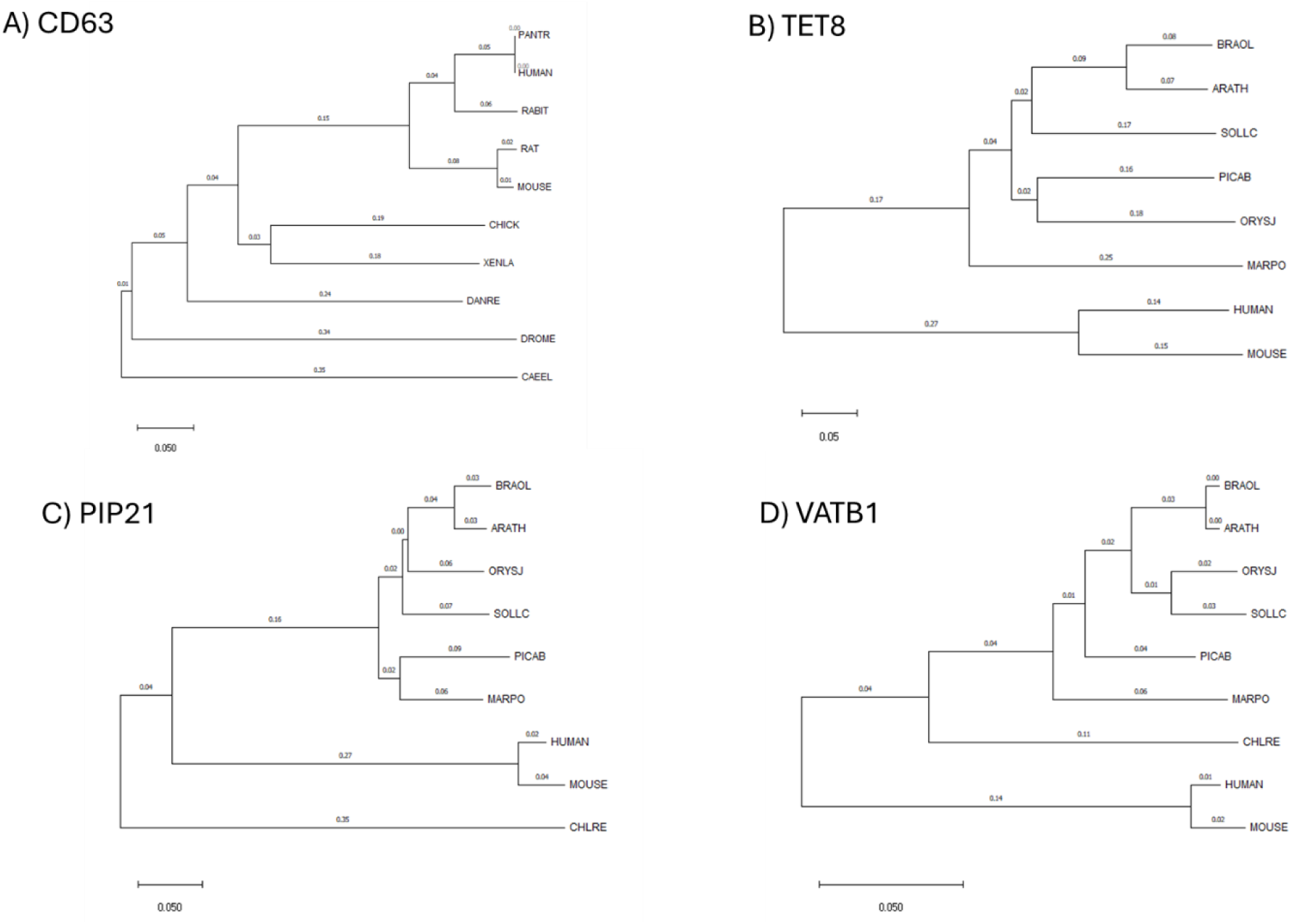
Phylogenetic trees. Evolutionary distance representation of A) CD63, B) TET8, C) PIP21, D) VATB1. The scale bar for potential PEV markers′ orthologues is set at 0.05. Phylogenetic trees were generated using MEGA11^23^.

Two proteins already defined as EV markers were included to serve as benchmarks, allowing us to determine whether the evolutionary distances of our proposed markers fall within acceptable ranges: CD63, a specific marker for animal species, and TET8, defined as a specific marker for PEVs and also detected in animals EVs. The other two proteins highlighted in **Figure 5** correspond to PIP21 and VATB1, both found to have high specificity for vesicles and of great abundance in our data.

The phylogenetic tree for CD63 (**Fig. 6A**), using *H. sapiens* as reference, showed that putative orthologues from mammalian species form a distinct cluster in the upper branches, with minimal evolutionary distances between them. This cluster was followed by a more distant grouping of proteins from amphibians, birds, and fish. At the most distal evolutionary branch, proteins from invertebrate species were observed. Notably, the total evolutionary distance between *H. sapiens* and the most distant species, *C. elegans*, was 0.69. CD9 and CD81 phylogenetic trees displayed a similar pattern (**Suppl. Fig. S4**).

For the TET8 phylogenetic tree (**Fig. 6.B**), as expected, the putative orthologues from species closely related to *A. thaliana*, such as other angiosperms and gymnosperms, exhibited the shortest evolutionary distances. No orthologue was found for chlorophytes, and orthologues from the animal kingdom appeared on the furthest branches, reflecting the largest evolutionary distances. The distance between *A. thaliana* and the most distant plant orthologue, TET8 from *M. polymorpha L.,* was 0.47, and in comparison, to *H. sapiens*, was 0.8.

The phylogenetic tree obtained for PIP21 demonstrated a higher conservation degree to that of TET8, with angiosperms and gymnosperms showing quite short evolutionary distances (**Fig. 6C**). Orthologues from non-plant species were grouped together but were clearly separated from the plant clade, showing a substantial evolutionary distance of 0.49. The chlorophyte *C. reinhardtii* (where no ortholog for TET8 was found) remained the most distant orthologue, with an evolutionary distance from *A. thaliana* of 0.61. Moreover, aquaporin 1-2 (PIP12) and clathrin heavy chain 2 (CLAH2) showed evolutionary distances comparable to those of PIP21, and notably shorter than those observed for TET8, along with a similar distribution pattern across phylogenies **(Suppl. Fig. 5)**. In contrast, syntaxins 121 (SY121) and 132 (SY132), as well as calreticulin-1 (CARL1), displayed evolutionary distances more similar to TET8 **(Suppl. Fig. 6 and 7A).** In some cases, however, the algae ortholog appeared more distant than the orthologous proteins found in animal species.

**Figure 7.**
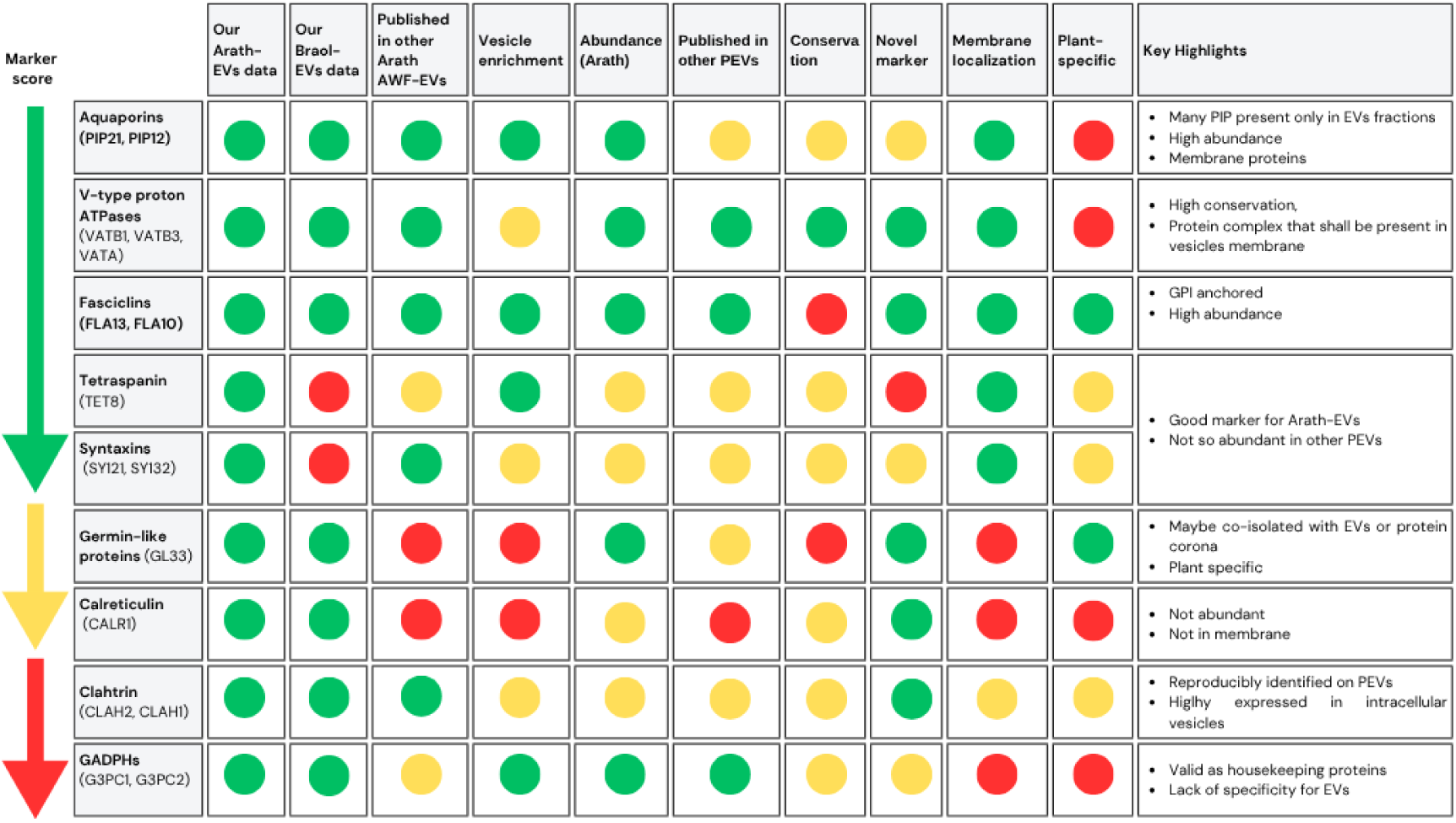
PEV marker score based on the criteria used to evaluate each protein. The table organizes proteins based on various evaluation criteria. Each column displays a color code (in order of greater value: green, orange, red) to reflect specific thresholds for each criterion. 1) Presence in our datasets of Arath-EVs: green = present; red = absent. 2) Presence in our datasets of Braol-EVs: green = present; red = absent. 3) Published in other AWF-Arath-EVs datasets: green = present in all 3; orange = 2; red = 1 or none. 4) Vesicle enrichment: green = present only in EVs fractions or high enrichment; orange = moderate enrichment; red = low enrichment 5) Abundance ranking (based on our data from Arath-EVs): green = Top 300; orange = 300 - 400; red = below 400. 4) 6) Presence in other PEVś datasets: green = present in 5 - 6; orange = 3 - 4; red = 1 - 2. 7) Conservation: green = very high; orange = high; red = acceptable (plant-specific). 8) Novel markers: green = novel markers we propose; orange= proposed in some studies; red = already used/proposed as markers: 9) Membrane localization: green = transmembrane or GPI-anchored; orange = potentially associated with the membrane (e.g., through lipid interactions); red = membrane localization unlikely. 10) Plant-specificity: green = no orthologues in animals; orange = orthologues with less than 30% of identity; red = orthologues found in animals (>30%).

V-type proton ATPase subunit B 1 (VATB1) displayed a similar evolutionary trend for the plant species than the TET8 tree, though with much shorter phylogenetic distances (**Fig. 6D**). The evolutionary distance between the orthologues of *A. thaliana* and *M. polymorpha* was 0.12, the shortest distance among all the proteins evaluated. Additionally, the evolutionary distance between *A. thaliana* and *H. sapiens* VATB1 was 0.28, supporting the high level of conservation across species and kingdoms. This conservation pattern, based on evolutionary distances, was similar to the observed for VATA **(Suppl. Fig. 7B)**

Finally, FLA13, FLA10, and germin-like 3 (GL33), which we propose as PEV-specific markers, showed larger evolutionary distances **(Suppl. Fig. 8 and 9).** These three proteins are plant-specific and, as expected, exhibited the highest evolutionary distances within the plant kingdom among all proteins analysed. Notably, the fasciclins FLA13 and FLA10 showed clearly greater distances than those observed for CD63. In contrast, GL33 displayed evolutionary distances more comparable to CD63, remaining within acceptable thresholds similar to those previously described for CD63 and TET8.

## Discussion

Although the field of plant EVs needs to fulfill the gap of marker determination to comply with the guidelines of the ISEV (MISEV), this has not been accomplished so far due to the lack of a compendium of *bona fide* markers of PEVs that could be used as a reference. In this work, we have paved the way for the standardization of PEV markers, assessing the utility of current and some novel potential markers based on an in-depth proteomic analysis of two reference plant species, as well as by conducting an exhaustive analysis of previous available EVs′ proteomic data from any plant species. Notably, the PEVs used in this study were isolated from apoplastic washing fluid after applying relevant controls (AWF dilution factor by indigo carmine and intracellular contamination by MDH activity assay) to Arabidopsis AWF as a proof-of-concept, therefore ensuring these are enriched in genuine extracellular vesicles. Regarding genuine EV isolation, several methods have been used to extract nanovesicles from plant tissues, such as blending, crushing or fruit juicing, which imply a certain degree of contamination with intracellular vesicles. However, the only approaches that reliably ensure a considerable enrichment of extracellular vesicles shall be: 1) the use of apoplastic washing fluid, 2) precise collection of culture medium from cell or tissue/callus-derived cells, and 3) collection of plant exudates, phloem and xylem sap. The former, used in this study, has the advantage of using whole tissue instead of cell cultured explants.

In order to overcome the absence of raw proteomic data available in repositories, we hereby provide the most profound proteomic data from apoplastic washing fluid-derived EVs from *A. thaliana.* and *B. oleracea* obtained so far. To accomplish this, we have optimized EV lysis and protein solubilisation methodologies to extract proteins from SEC-isolated EVs for LC-MS/MS analysis, based on a two-step technique. Proteomic analysis of the EV lysis supernatant, compared to that of the solubilized pellet, demonstrated that most proteins from the first extract (94 out of 117, **Suppl. Table 1 and Suppl. Fig 1B**) were as well identified in the second extract, which shall additionally be enriched in membrane bound proteins, that may potentially contain more relevant EV markers. Besides, among the 23 proteins exclusively identified in the EV lysis sample **(Suppl. Table 1C)** we could not find any potential EV marker. For these reasons, we focused on the second extract, identifying 1145 proteins in *A. thaliana* and 899 in *B. oleracea*, detected in all 3 biological replicates assayed (**Suppl. Table 2, Fig. 2E and 2F**). Next, we compared both extracts, searching for orthologues identified in both species with potential as EV markers, focusing on proteins related to EV biology, arising from the bioinformatic post-analysis (**Fig. 3**). Twelve protein families were found consistently identified in both EV proteomes: fasciclin-like arabinogalactan proteins, germin-like proteins, patellins, heat shock proteins, aquaporins, calreticulins, clathrins, phospholipase Ds, Rab proteins, V-type proton ATPase subunits, 14-3-3 like proteins and GAPDHs (**Table 2**). All the protein orthologues showed more than 75% of identity, most being above 90%. Since *A. thaliana* and *B. oleracea* species belong to the *Brassicaceae* family of angiosperm eudicot plants and therefore, our analysis shall rule out species-specific markers, since those proteins not reproducibly identified in plant material from the same family would rarely be useful to identify EVs from other subdivisions of the plant kingdom. Interestingly, none of the classical Arath-EVs PEVs markers were consistently identified in both proteomes: TET8 is present in *A. thaliana,* but only detected in one replicate of *B. oleracea*; TET9 was not identified in any replicate of both; and syntaxin 121 (SY121 or PEN1) present in *A. thaliana* but not detected in any replicate of *B. oleracea.* Besides, EXO70E2 was not detected in any EV extract, which may imply a low contribution of EXPO-derived EVs to the AWF-EV pool. Concerning orthologues of mammalian EV markers suggested in literature^6^, TSG101 and Alix were not detected in any of the replicates from both plant EVs. The absence of reproducible identification of these markers in Arath-EVs may be explained by a low protein abundance. This shall be enough reason to consider these as unideal markers for species other than *Arabidopsis*, in favor of those that we could consistently identify, since their determination may be complicated due to their low abundance. Anyhow, we can never rule out the possibility that all these proteins′ peptides may not be the most favorable to be detected by LC-MS/MS, which could explain why they were not identified. Furthermore, antibody-based assays may allow identifying some of these markers even at low abundance, but this will not guarantee that the marker is present in a considerable number of EVs from the sample obtained.

Since *A. thaliana.* is the best annotated species in UniProt database (the one used in our proteomic analysis) and has the most PEV proteomic studies published to date, we selected the three reference datasets from AWF-derived Arath-EVs published so far^3,24,25^ to compare them with our results. Considering that raw data was not deposited in repositories in the three cases, we compared the list of identified proteins in all 4 studies, finding 34 proteins in common **(Fig. 4C)**. Among these, we found fasciclin-like arabinogalactan proteins 2 (FLA2) and 8 (FLA8), heat shock 70 kDa protein 1 (HS701), aquaporins PIP1-2 and PIP2-7, clathrin heavy-chain 1 (CLAH1) and 2 (CLAH2), phospholipase D alpha 1 (PLDA1), Ras-related protein RABE1c (RAE1C), V-type proton ATPase subunits A, D, D1, E1 and H; 14-3-3-like protein GF14 epsilon (14310) and syntaxin 121 (SY121 or PEN1), 71 (SYP71) and 132 (SY132).

In order for these potential EV markers to be considered valid identity markers, their specificity for differentiating EVs also needs to be considered. EV preparations are not completely pure and contamination with a certain degree of protein aggregates has to be assumed. In fact, post-analysis of the protein content from Arath-EVs and Braol-EVs shows presence of ribosomal, cytosolic and chloroplastic proteins **(Fig. 3A and Suppl. Fig. 3)**. Although some of the GO term annotations may not be accurate, since many proteins can be present in different cellular compartments, the co-isolation of some unspecific proteins in our PEV preparations can be envisioned. With the goal in mind of defining the specificity of the potential EV markers studied, we analysed the protein content of the SEC fractions not pooled for isolating EVs and calculated an enrichment degree in EVs, as the ratio = LFQ in EVs / LFQ in soluble protein fractions. This analysis offered key results to determine marker specificities and pointed out to some of the potential EV markers being equally present in EVs and the soluble AWF proteome or even enriched in the latter. In particular, patellin 1, hsp70, 14-3-3 proteins (9 out of 10) and the fasciclins FLA1, FLA2, FLA7 and FLA8; have not shown specificity for PEVs. On the contrary, TET8 and all the syntaxins analysed, including PEN1, demonstrated to be enriched in PEVs. Of note is the result obtained with the aquaporins, all of which were highly enriched in PEVs (11 of them only found in PEVs and not detected in the soluble AWF proteome, and PIP21 found with a 23.2-fold enrichment). Furthermore, most V-type proton ATPase subunits (7 out of 9) appear enriched in PEVs. Although some fasciclins have been shown to have low specificity for EVs, FLA10 and FLA13, were enriched more than 10 times in PEVs, and FLA18 was found enriched more than 5 times. As for the case of the germin-like proteins, 2 out of the 3 analysed (GL31 and GL33) were found enriched in PEVs. All GAPDHs, calreticulins and clathrins analysed were found to be PEV-specific.

Aiming to investigate the potential of the PEV-specific markers found, we searched for the appearance of these protein families in previous proteomic studies of EVs from other plant sources. The proteomic datasets selected had to fulfill the following criteria: 1) Isolating genuine extracellular vesicles (intracellular vesicles underrepresented); 2) Depth of > 100 proteins identified; 3) Valid isolation methods according to MISEV guidelines. Concerning the first criterium, we have included fruit juice-derived EVs as a mild homogenization method providing an isolate enriched in EVs, since the potential contribution of intracellular nanovesicles to these extracts is expected to be minimized, but conclusions extracted from such studies may not be as specific as those from AWF, or else pollen or tissue/callus/cell culture supernatants^2^. All families studied but germin-like proteins and calreticulins were found in at least 50% of the PEVś datasets analysed, supporting their recurrent identification in EVs from other plant sources. FLA18 was not found in any other study and, since it is not a GPI-anchored fasciciclin, we have decided not to prioritize it as a robust PEV marker. This reinforces their utility as markers for PEVs but also evidences the difficulties that we may be facing aiming to find EV protein markers valid across different plant species, due to probable species-specific protein expression patterns and the fact that different EV subtypes shall contain distinct protein markers. Among the markers that showed specificity for PEVs, clathrin heavy chains 1 and 2 are reproducibly identified on PEVs but may not be ideal markers since they are highly expressed in intracellular vesicles. Concerning GAPDHs, their consistent identification may support their use as housekeeping proteins in certain assays, but we can anticipate a lack of specificity as PEV markers due to their ubiquitous expression in cells and tissues.

On the other hand, V-type proton ATPase subunits A, B, Dand H arise as novel EV markers, all belonging to a vacuolar ATPase protein complex that shall be present in the membrane of these vesicles. This protein complex has not been either proposed as EV marker in mammalian cells, but its presence in EVs from several animal species has been reported in Vesiclepedia^30^ (**Suppl. Table 6**). For these reasons, we believe that the V-type proton ATPase represents a novel EV marker with great specificity both for PEVs and animal EVs.

Among the protein families with potential to be used as PEVs markers, there are two that are plant-specific: germin-like proteins^31^ and fasciclin-like arabinogalactan proteins (FLAs)^32^. There are several advantages of using plant-specific markers to identify EVs: 1) Ruling out cross-contamination with animal material; 2) Unequivocally targeting PEVs on animal *in vitro* and *in vivo* settings; and 3) Possibility to differentiate PEVs from EVs from bacteria/fungi/oomycetes in host/pathogen and plant/microbe interaction studies. Germin-like proteins are extracellular matrix proteins belonging to the apoplast^31^. For this reason, although their enrichment in PEVs has been hereby proven, further analyses will be required to confirm their presence in PEVs, since they could either be encapsulated within PEVs or be a part of the protein corona. The case of the FLAs deserves particular attention. These proteins are plant-specific and while some of the FLAs found were also found in the soluble fraction of the AWF, the two found to be EV-specific, FLA10 and FLA13, are unequivocally bound to the plasma membrane through a GPI-anchor^33^. Furthermore, there are no other variants described for these proteins that could be secreted instead of anchored to the plasma membrane^32^. GPI-anchored proteins are included in MISEV2023^1^, together with transmembrane proteins, in the first class of protein markers considered EV hallmarks, therefore as *bona fide* markers of EVs. Proteins from this family have a common protein domain, FAS1^32^, a cell adhesion domain conserved in plant and animals. Nonetheless, animal proteins containing this domain lack arabinogalactan polysaccharides and a GPI-anchor, which implies alternative subcellular localization and results in completely different functions^34^.

As a result of all the analyses performed, we chose certain candidates among the protein families found to show specificity for PEVs to further analyse the presence of orthologues and their degree of conservation throughout the plant kingdom: TET8, PIP21, PIP12, VATB1, VATA, CLAH2, SY121 (PEN1), CALR1, FLA13, FLA10 andGL33.. Besides, representative species of the different plant taxonomy subdivisions, *Homo sapiens* (human) and *Mus musculus* (mouse) species were added to the analysis to evaluate the global degree of conservation of the analysed markers. Likewise, in order to compare the degree of conservation of these PEV markers with that of the current EVs markers used in animal-derived EVs, the most used: CD63, CD81 and CD9^1^, were chosen. The results obtained for VATB1 and VATA point to a high degree of conservation throughout plants and animals. This may imply functional conservation and support their use as markers for both animal EVs and PEVs, with the drawbacks of not ruling out cross-contamination of EVs from other sources and the impossibility to differentiate EVs from different origins when necessary. A subset of proteins (CLAH2 and aquaporins PIP12 and PIP21) do have orthologues in human and mouse, but these display low homology with those of plants, while all orthologues throughout the plant kingdom show short phylogenetic distance in all species but the algae *C. reinhardtii*. For the rest of the proteins analysed, *C. reinhardtii* shows the greatest phylogenetic distance among all plant species and is very close to that of human and mouse orthologues, when these exist. Furthermore, we could not find an annotated ortholog in *C. reinhardtii* for GL33. This seems to imply that most of the hereby proposed PEVs markers may not be applicable to algae-derived EVs or at least may need further confirmation in such samples. TET8 and PEN1 (SY121) showed greater evolutionary distances to those of v-type ATPases, aquaporins and clahtrins, and similar to CALR1 and the other syntaxin SY132. The latteŕs ortholog in algae was far more distant than that of PEN1, even above the distance found for the human and mouse orthologues.

Three proteins were confirmed to be plant specific: FLA10, FLA13 and GL33, since no orthologues in human nor mouse were found. Their degree of conservation in plants was compared with the degree of conservation of classical markers used in mammalian cell-derived EVs in the animal kingdom. The phylogenetic distances among the same plant family (*Brassicacea*) were analogous to that of human to rabbit and rodent orthologues. The distance of the mammalian markers from human to bird, fish and amphibia orthologues is similar to that of *A. thaliana* GL33 to other phanerogams: such as another eudicot family member (*Solanum lycopersicum*) and gymnosperms (*Picea abies*). We could not find an ortholog of GL33 in *C. reinhardtii*, as aforementioned, but the distance with the liverwort *M. polymorfa* is also analogous. On the contrary, human CD63, CD81 and CD9 have a much shorter distance to invertebrates *(Drosophila melanogaster* and *Caenorhabditis elegans*) than that of FLA10 and FLA13 with cryptogams (*bryophyta* or mosses and liverworts; and *cryptophyta* or algae). These data suggest that fasciclins FLA10 and FLA13 possess a lower degree of conservation in the plant kingdom than the most currently used markers in the field of EVs have throughout animal species.

To sum up, we provide **Figure 7**, an overview of the key aspects considered in this work to evaluate these protein families as potential PEV markers. Although integrating all these factors is challenging, we have categorized the families into three groups based on our criteria. Proteins ranked at the top are those we believe possess the most favorable characteristics for serving as vesicle markers. Additionally, noteworthy features for each protein are highlighted.

## Conclusions

Altogether, the results hereby presented offer a wide list of potential markers for genuine extracellular vesicles derived from plant sources, providing evidence of EV specificity, identification in previous proteomic studies, as well as the existence of orthologues and conservation throughout the plant kingdom, together with human and mouse species, as reference animal proteomes. The data gathered shows consistent PEV specificity for the current PEV markers used in *Arabidopsis thaliana* TET8 and PEN1, but these are not robustly identified in our proteomic data from *A. thaliana* and *B. oleracea*, and in some of the other PEV proteomes published so far. Hence, alternative markers need to be explored aiming to create a broader set of standard PEV identity markers available for the scientific community. Taking into account the specificity for EVs and abundance in the proteomic results of Arath-EVs in the present study, as well as the consistency of their presence across diverse datasets of PEVs from different species, we provide evidence supporting the potential of the following protein families as additional PEV markers: aquaporins, vacuolar-type ATPase complex subunits, fasciclin-like arabinogalactan proteins FLA 10 and FLA13, syntaxins other than PEN1, germin-like proteins and calreticulins.

## Acknowledgements

We would like to thank Prof. Antonio Marcilla for proofreading the manuscript.

## Funding received

*Programa de Atracción de Talento (Modalidad 1), Comunidad Autónoma de Madrid 2023-5AIND-28938* and *Ministerio de Ciencia, e Innovación y Universidades y Fondo Europeo de Desarrollo Regional PID2021-126274OB-I00; CNS2022-135368*. Dr. Ibone Rubio was supported by *Ayudas para la contratación de ayudantes de investigación y técnicos de laboratorio, Comunidad Autónoma de Madrid: PEJ-2023-AI/SAL-GL-27307*. Miriam M. Rodríguez de Lope was supported by *Ayudas para la realización del Programa Investigo, Comunidad Autónoma de Madrid (INV/09-PIN1-00013.4/2022)*.

