## Supplementary material for "A compendium of *bona fide* reference markers for genuine plant extracellular vesicles and their degree of phylogenetic conservation": Suppl Figures 1-9.pptx

### Slide 1
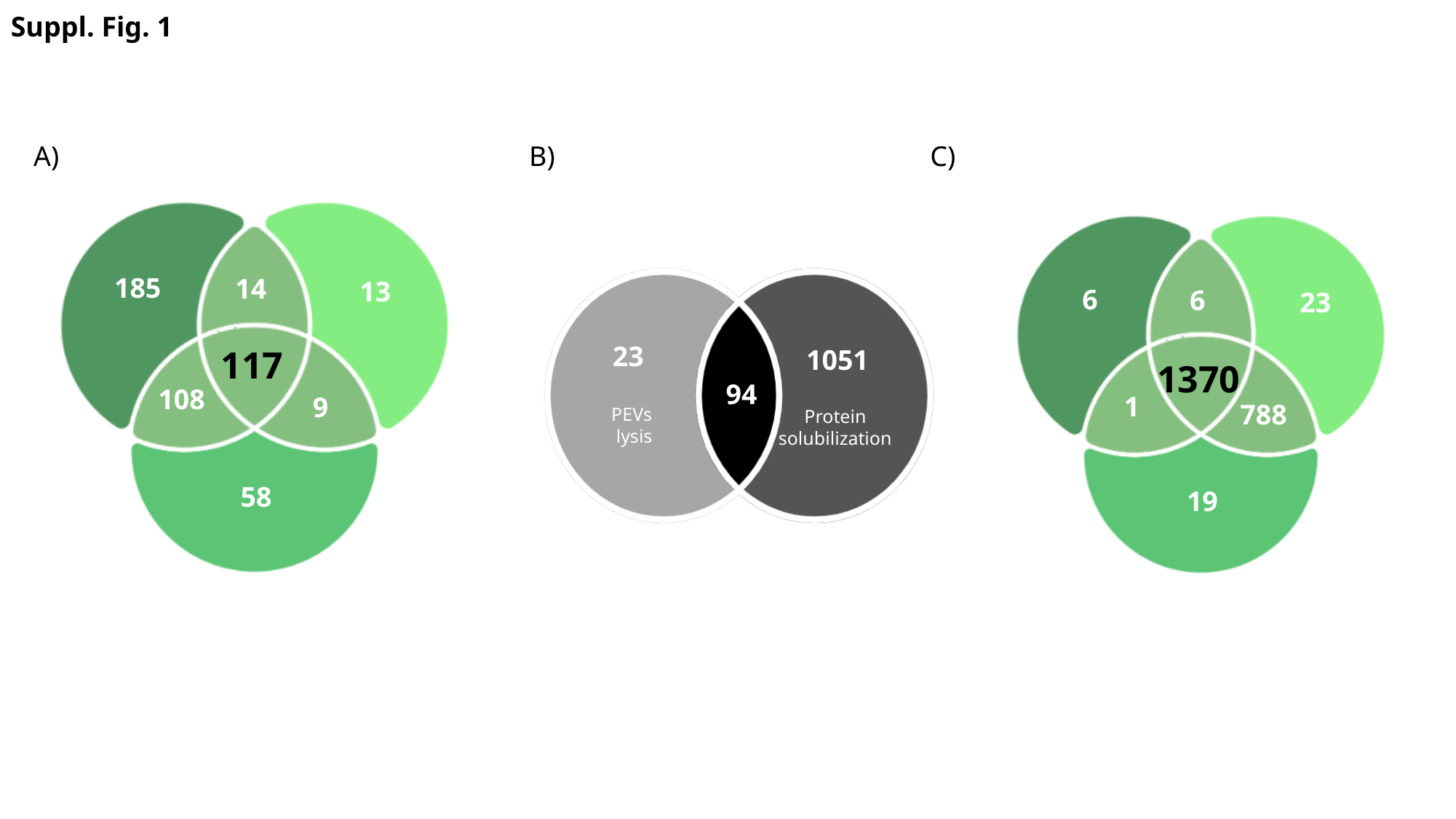

Suppl. Fig. 1
A)
B)
C)
185
14
13
117
108
9
58
6
6
23
1370
1
788
19
23
1051
94
PEVs
lysis
Protein
solubilization

### Slide 2
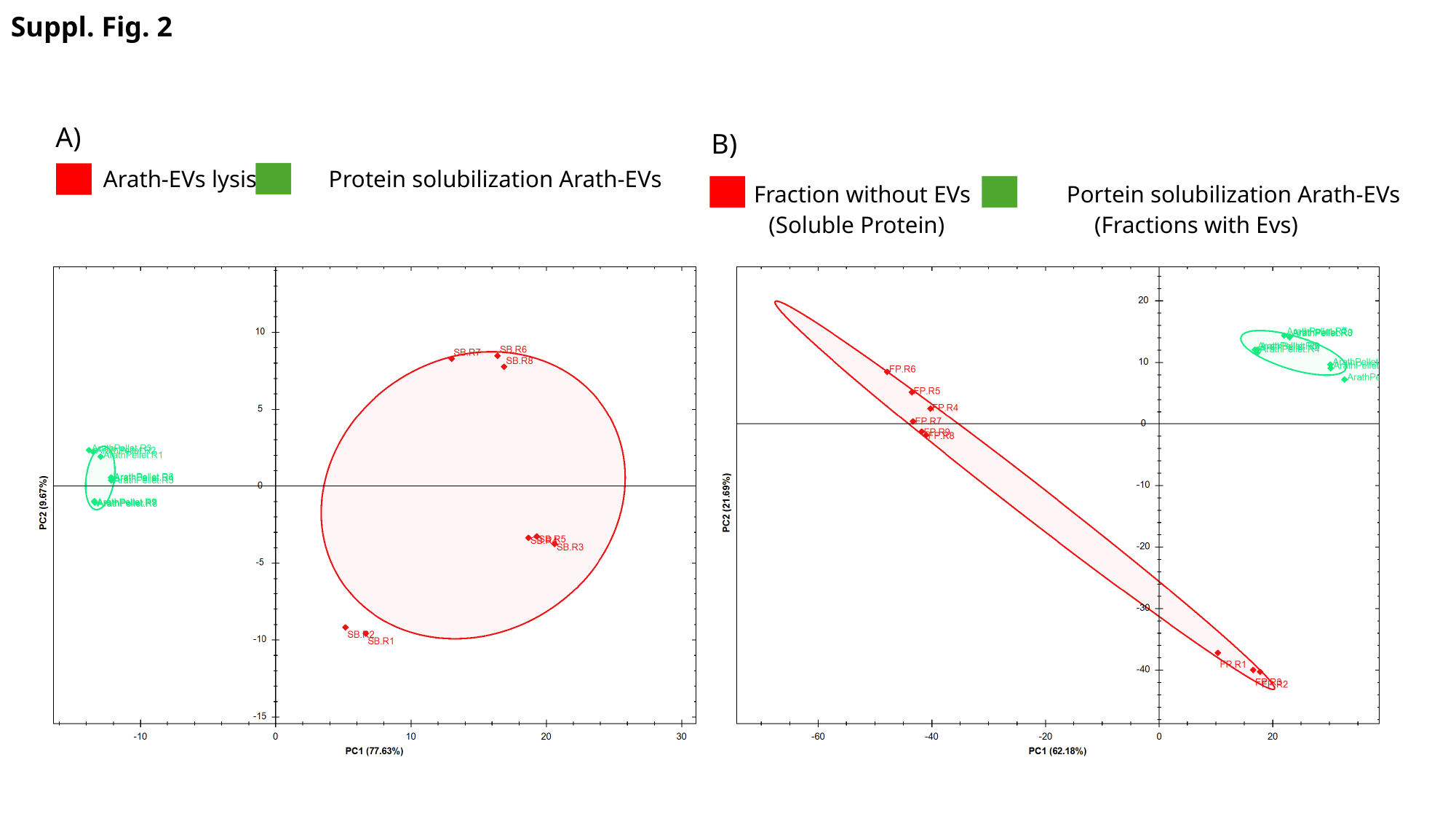

Suppl. Fig. 2
A)
B)
Arath-EVs lysis Protein solubilization Arath-EVs
Fraction without EVs Portein solubilization Arath-EVs
(Soluble Protein) (Fractions with Evs)

### Slide 3
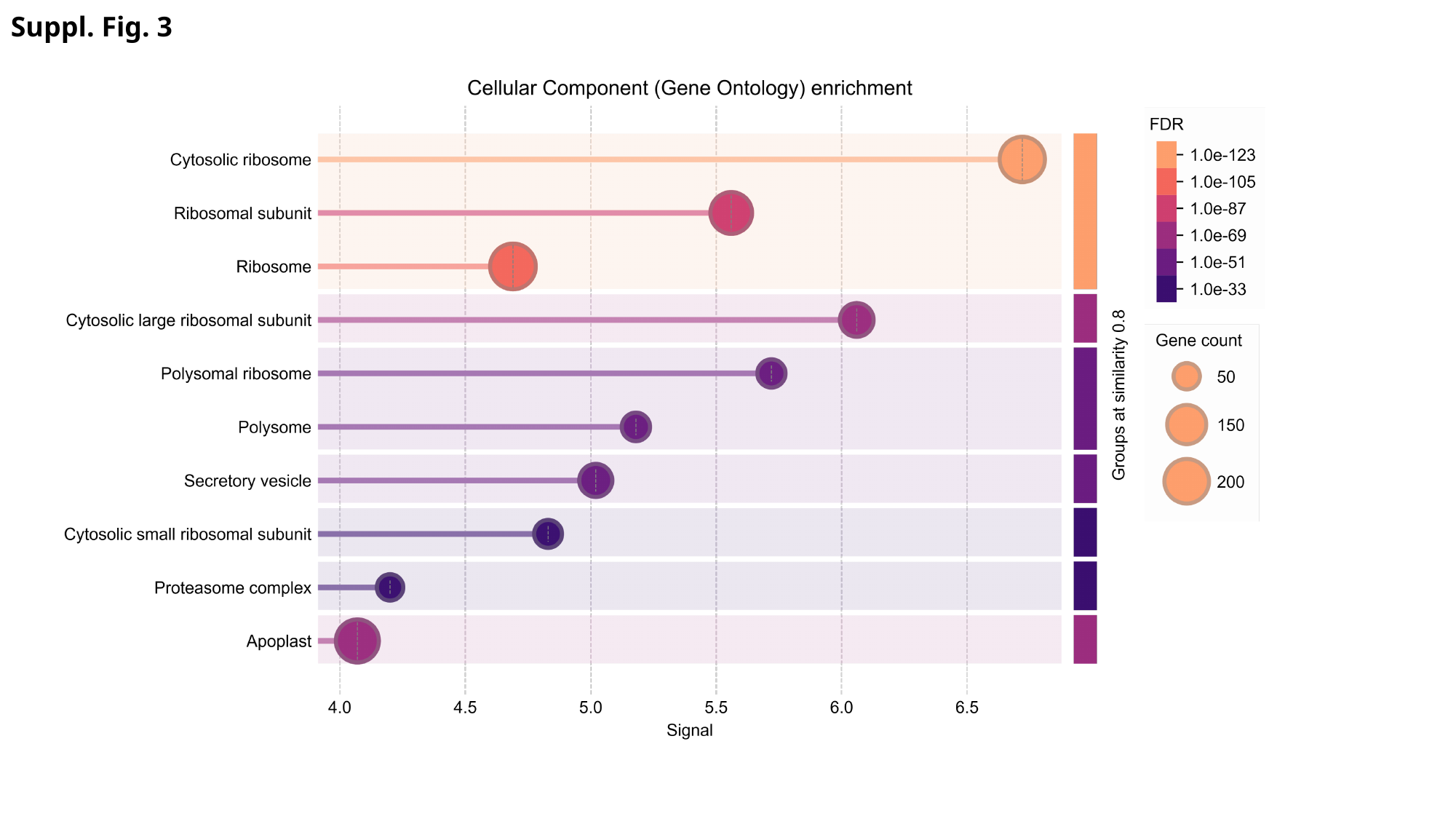

Suppl. Fig. 3

### Slide 4
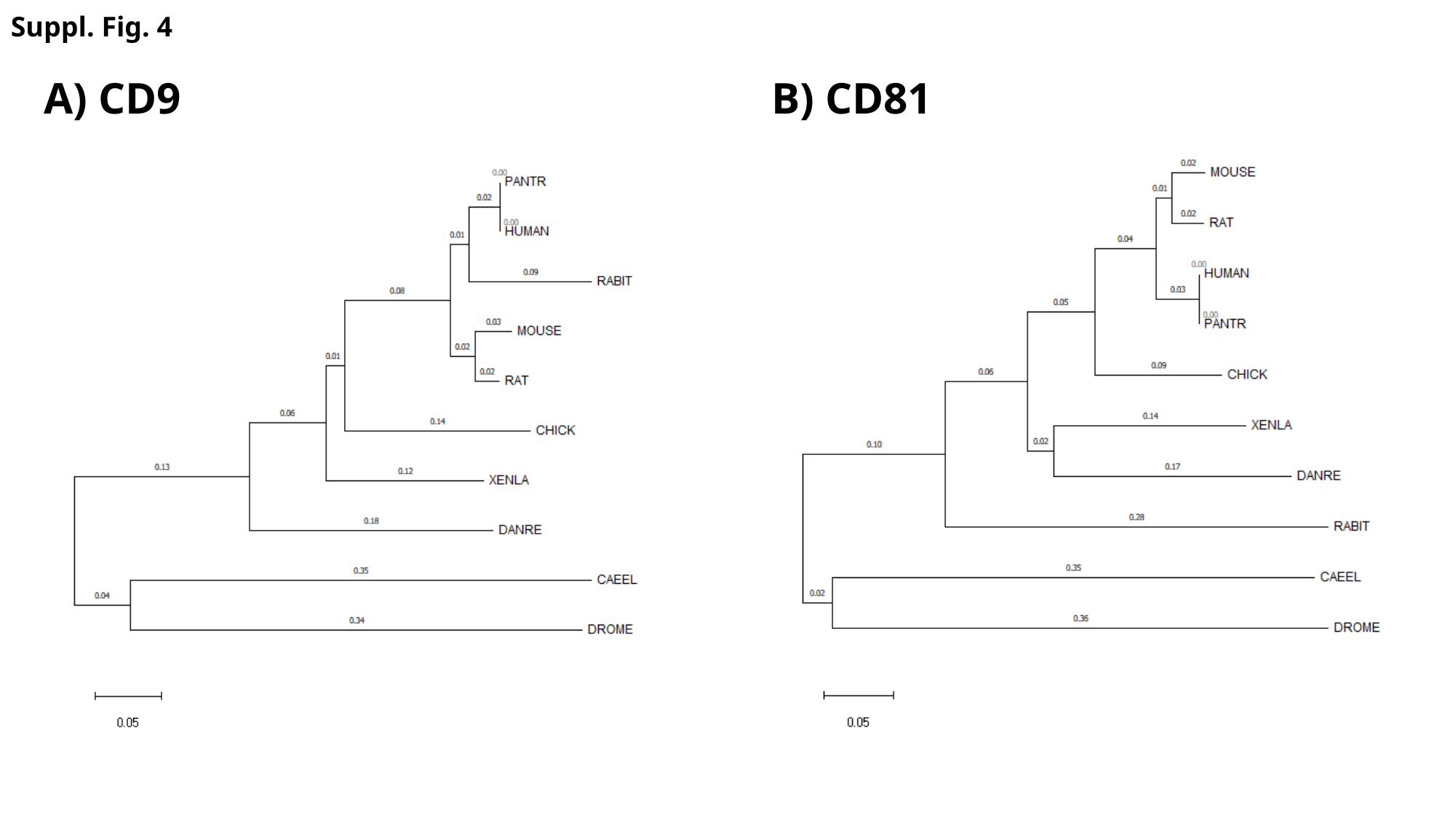

Suppl. Fig. 4
A) CD9
B) CD81

### Slide 5
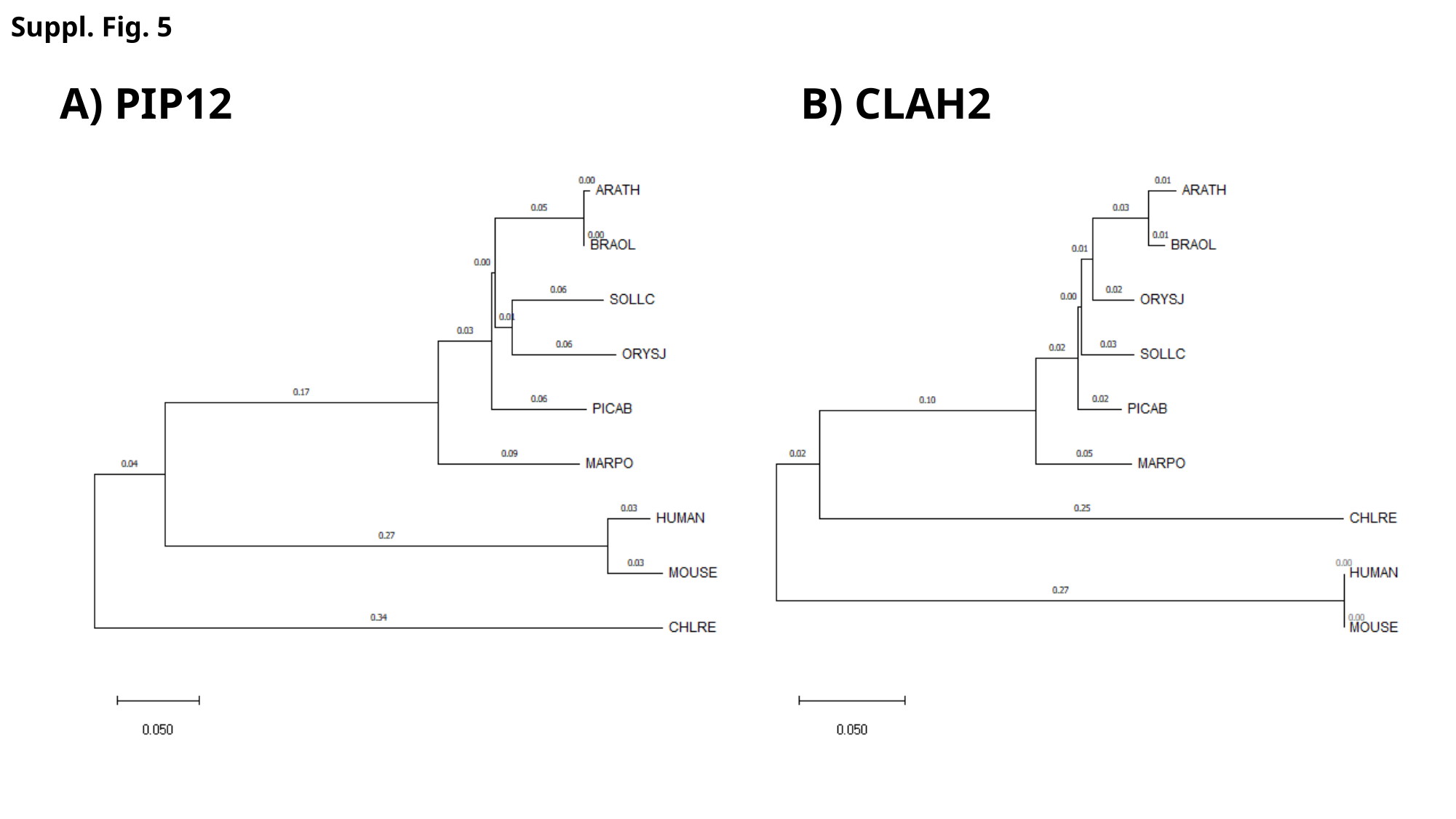

Suppl. Fig. 5
A) PIP12
B) CLAH2

### Slide 6
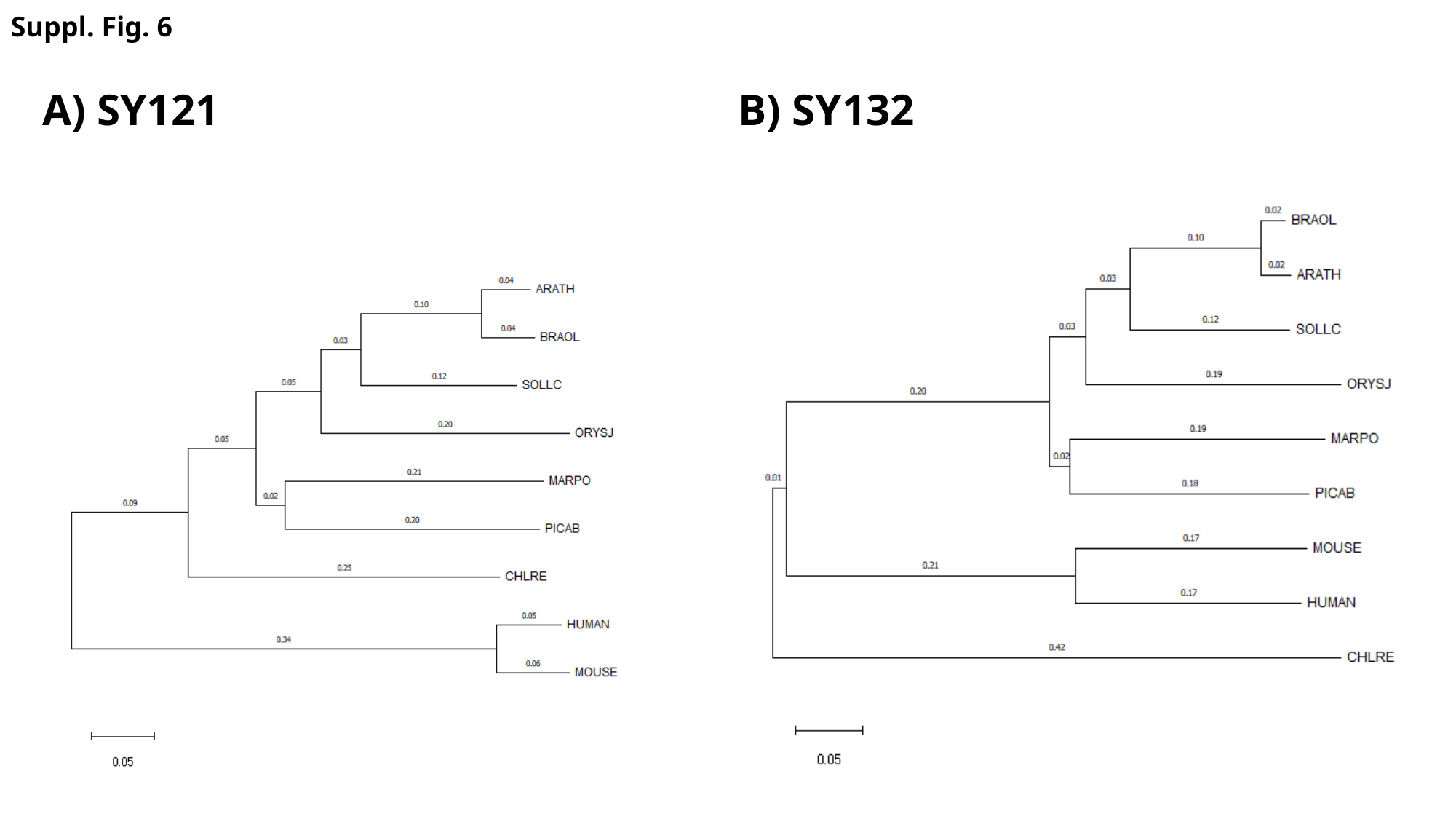

Suppl. Fig. 6
A) SY121
B) SY132

### Slide 7
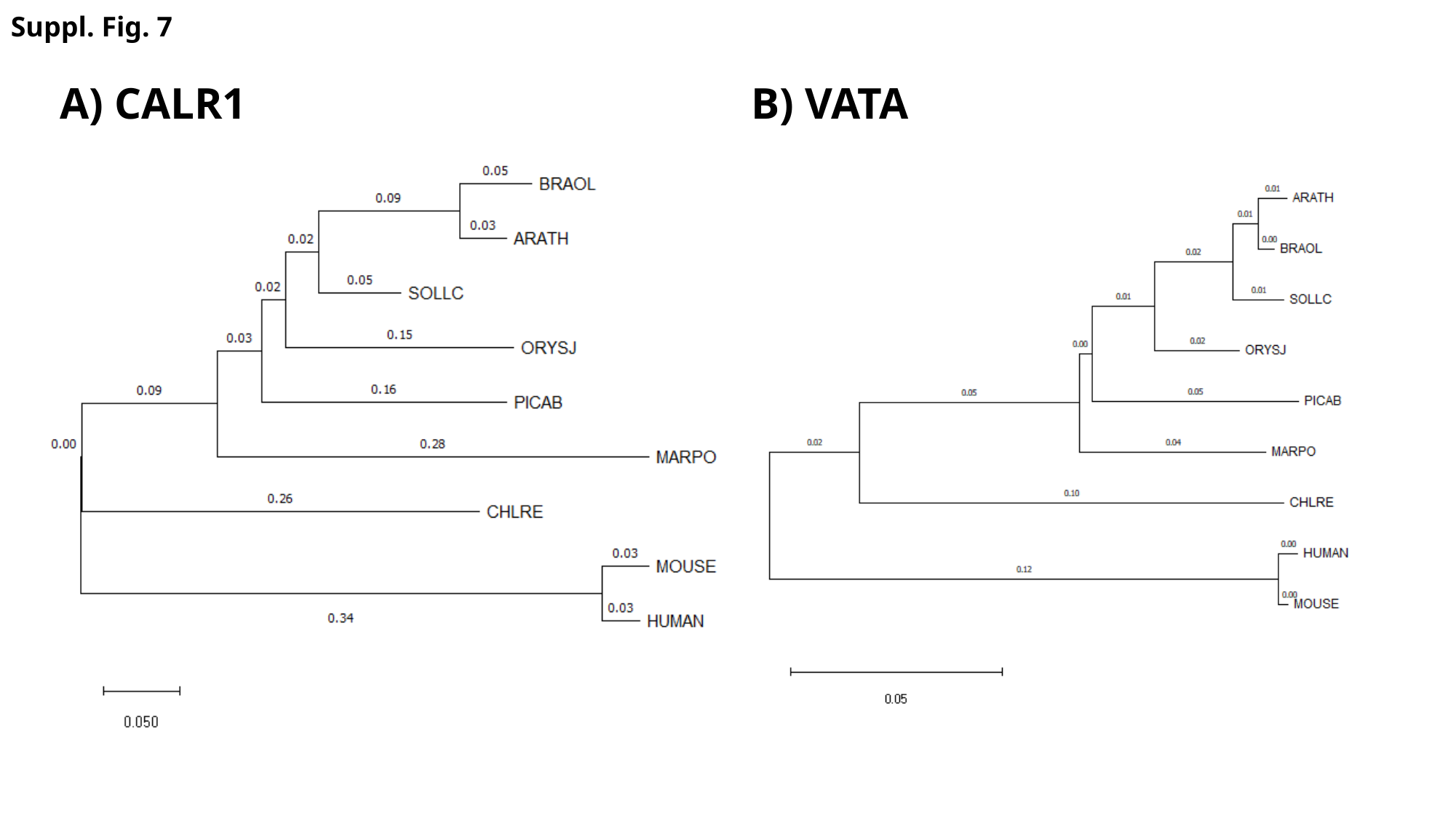

Suppl. Fig. 7
A) CALR1
B) VATA

### Slide 8
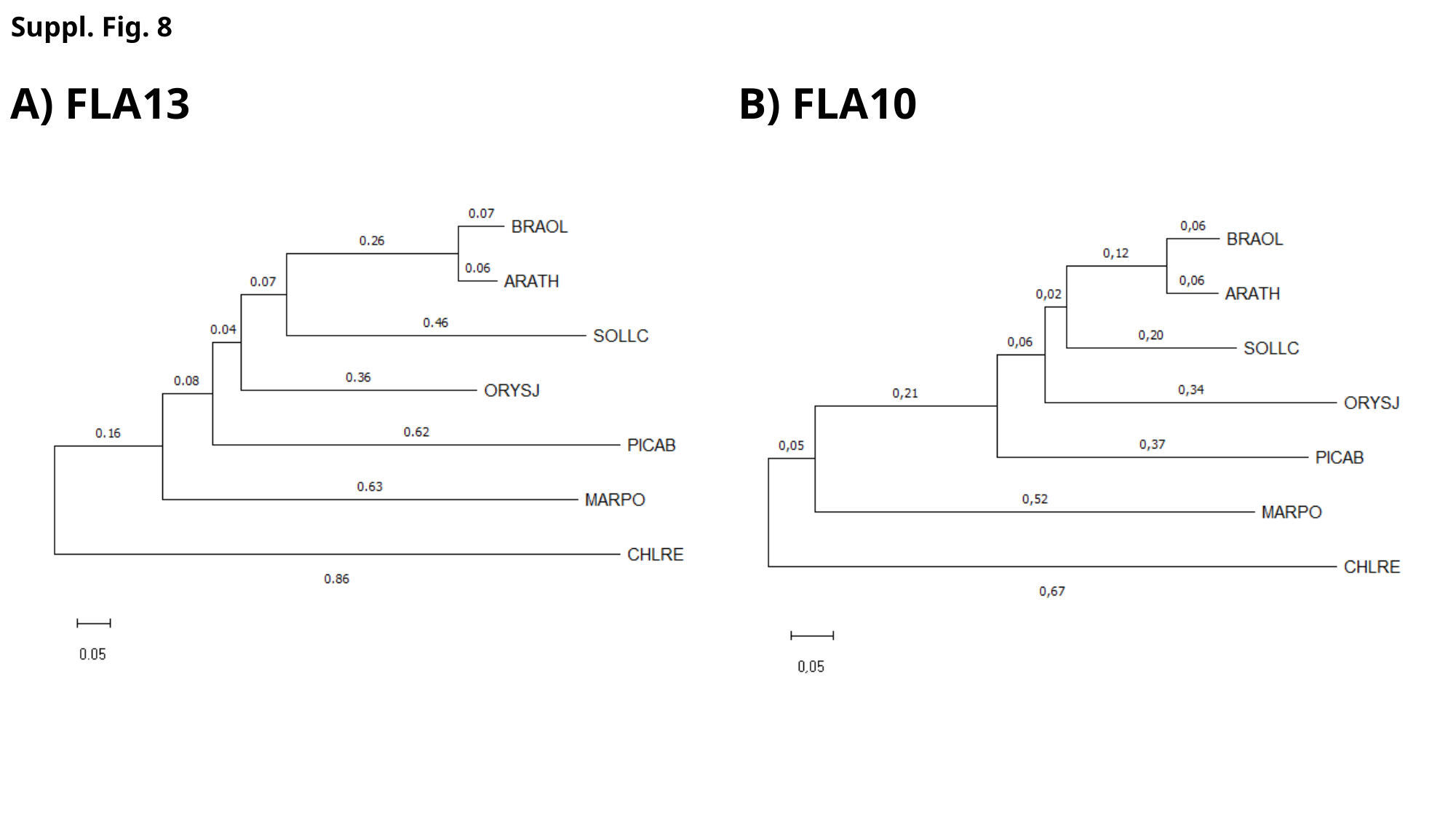

Suppl. Fig. 8
A) FLA13
B) FLA10

### Slide 9
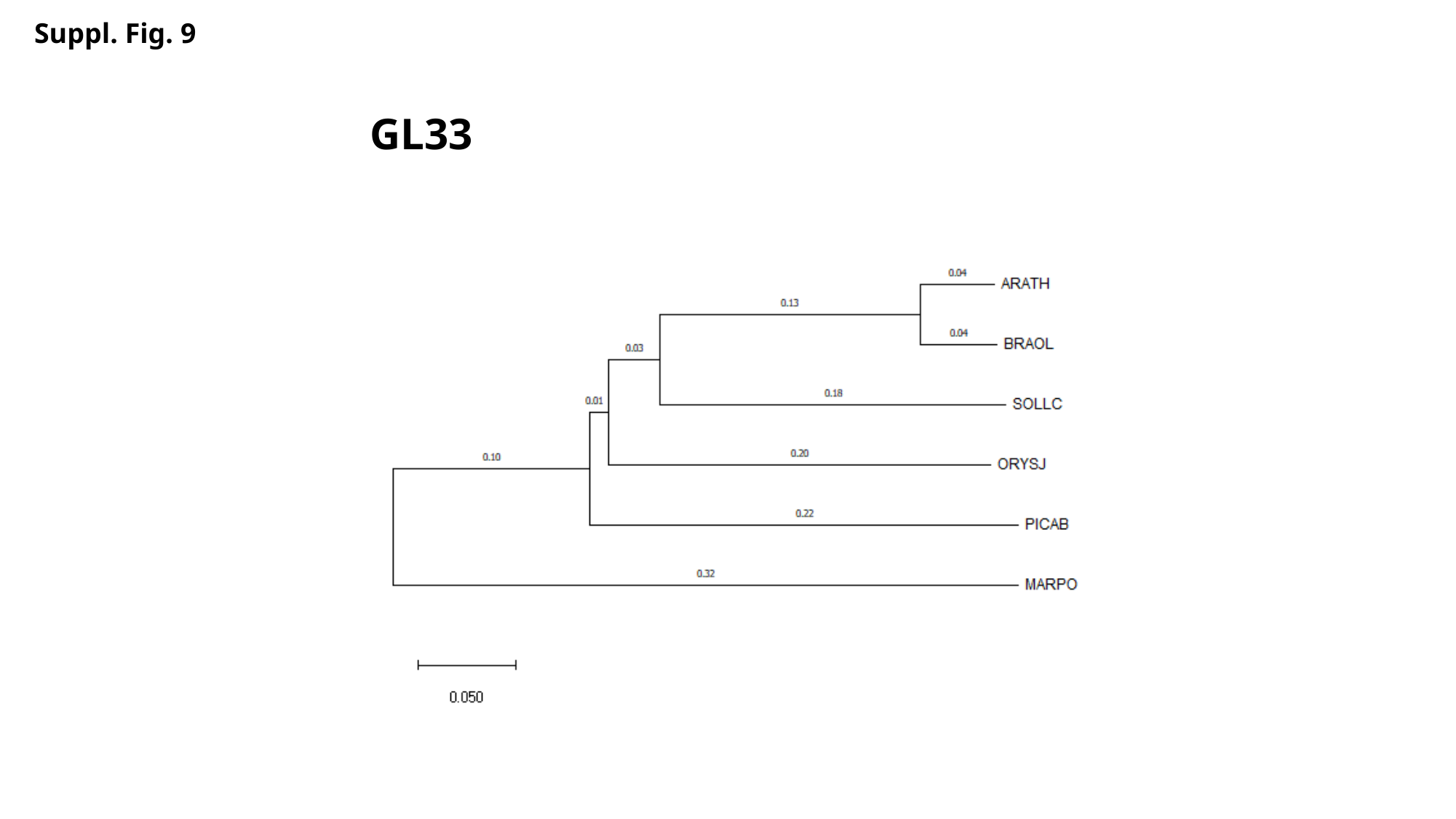

Suppl. Fig. 9
GL33
